# Persistent DNA damage rewires lipid metabolism and promotes histone hyperacetylation via MYS-1/Tip60

**DOI:** 10.1101/2021.06.24.449832

**Authors:** Shruthi Hamsanathan, Tamil Anthonymuthu, Steven J. Mullett, Stacy G. Wendell, Suhao Han, Himaly Shinglot, Ella Siefken, Austin Sims, Payel Sen, Hannah L Pepper, Nathaniel W. Snyder, Hulya Bayir, Valerian Kagan, Aditi U. Gurkar

## Abstract

Nuclear DNA damage is intricately linked to cellular metabolism. However, the underlying mechanisms and full range of metabolic alterations that occur in response to persistent DNA damage are not well understood. Here, we use a DNA repair-deficient model of ERCC1-XPF in *Caenorhabditis elegans (C. elegans)*, that accumulates physiologically relevant, endogenous DNA damage, to gain molecular insights on how persistent genotoxic stress drives biological aging. Using an integrated multi-omic approach, we discover that persistent genotoxic stress rewires lipid metabolism. In particular, nuclear DNA damage promotes mitochondrial β-oxidation and leads to a global loss of fat depots. This metabolic shift to β-oxidation generates acetyl-CoA and drives histone hyperacetylation. Concomitantly, we observe an associated change in gene expression of immune-effector and cytochrome (CYP) genes. We identify MYS-1, the ortholog of mammalian histone acetyltransferase TIP60, as a critical regulator of this metabolic-epigenetic axis. Moreover, we show that in response to persistent DNA damage, polyunsaturated fatty acids (PUFAs), especially arachidonic acid (AA) and AA-related lipid mediators are elevated. This elevation of PUFA species requires *mys-1*/Tip60. Together, these findings reveal that persistent nuclear DNA damage alters the metabolic-epigenetic axis to drive an immune-like response that can promote age-associated decline.

## INTRODUCTION

Persistent DNA damage not only promotes developmental disorders, but also drives a range of age-related chronic conditions like metabolic syndrome, cardiovascular disease, cancer and neurodegeneration^1,2^. Technological advances have strengthened the evidence for accumulation of DNA damaging lesions and DNA mutations with age and disease in several model organisms, as well as humans^1,3-6^. In humans, several inborn mutations of genes linked to DNA repair are associated with progeroid syndromes^1,7,8^. Furthermore, although pediatric cancer patients treated with DNA damaging chemotherapeutic drugs or radiotherapy have increased survival rates, they often exhibit frailty and develop multi-organ aging pathologies in their mid-40s^9-11^. Given that DNA damage occurs continuously throughout life, discerning the underlying mechanism of how genotoxic stress contributes to health and disease is critical.

Nuclear DNA repair is intimately tied to metabolism. In response to acute nuclear genotoxic stress, metabolic pathways regulate the availability of metabolite pools, which are essential for the DNA repair process^12,13^. In contrast, the full range of metabolic alterations that occur in response to persistent DNA damage are not well understood. Interestingly, certain interventions that target the metabolic node can alleviate persistent DNA damage-driven age-associated pathologies. For example, in *Ercc1*^*-/Δ*^ mice, a model where persistent DNA damage occurs due to loss of a DNA repair endonuclease, dietary restriction (DR) significantly improved healthspan and extended lifespan^14^. In *Csb*^*-/-*^ mice and in *C. elegans* rendered deficient in transcription-coupled nucleotide excision repair (TC-NER), a model of Cockayne syndrome, supplementation with a NAD+ precursor reversed metabolic, mitochondrial, and transcriptional changes, and improved overall health^15,16^.

Recent evidence has uncovered cellular metabolism as a modifier of the epigenome, providing support for the metabolic-epigenetic axis. For example, β-oxidation is the multi-step mitochondrial process of breaking down long chain fatty acid into units of acetyl-CoA. Acetyl-CoA is critical for histone acetylation and gene expression^17^. Some studies have reported changes to the epigenome in response to acute genotoxic stress^12^. Similarly, there is evidence that suggests that DNA damage may reprogram aspects of cellular metabolism^7,18,19^. However, whether the metabolic-epigenetic axis is altered in response to persistent DNA damage and whether these changes have an impact on organismal health is currently unknown.

ERCC1 (excision repair cross complementing-group 1) is a DNA repair protein that forms a heterodimer with XPF (xeroderma pigmentosum group F-complementing protein) and functions as a 5′-3′ structure-specific endonuclease. ERCC1-XPF is required for multiple nuclear DNA repair mechanisms that include, nucleotide excision repair (NER), DNA interstrand cross-link (ICL) repair and double strand break (DSB) repair^20^. Because ERCC1-XPF is involved in multiple DNA repair pathways, patients lacking this complex, exhibit phenotypically pleiotropic syndromes including XFE syndrome, Cockayne syndrome, and cerebro-oculo-facial skeletal (COFS) syndrome or cancer prone disorders such as xeroderma pigmentosum^1,7,21^. Like XFE progeroid patients, loss of ERCC1-XPF in mouse models, spontaneously leads to an accelerated aging phenotype that includes progressive sarcopenia, osteoporosis, cardiovascular disease, neurodegeneration and other well-known age-related morbidities^22,23^. To examine the complex effect of physiologically relevant, persistent endogenous DNA damage on health, we used *C. elegans* as a model system. Phenotypes associated with loss of ERCC1-XPF in mammals is recapitulated in *C. elegans*, including increased susceptibility to DNA damaging agents, accumulation of endogenous DNA lesions and premature aging.

We have previously demonstrated that it is not the DNA damage per se that drives the onset of disease, but rather the DNA damage-induced cellular responses that ultimately promote a decline in health^24^. In particular, there is an initial genotoxic stress-adaptive response, during which ERCC1-XPF deficient *C. elegans* and mice do not display any functional decline, and in fact are resistant to a number of stressors^24,25^. However, with persistent DNA damage, there is an *exhaustion* of this response leading to augmented stress sensitivity and physiological deterioration^24,26^. A large gap in our knowledge is the mechanistic understanding of the cellular and molecular alterations that happen during this *exhaustive* phase. The identification of these molecular alterations are critical for the rational development of strategies to promote health and homeostasis.

## RESULTS

### Persistent DNA damage reprograms lipid metabolism

To gain a comprehensive understanding of the cellular and metabolic responses to genotoxic stress during the ‘*exhaustive*’ phase, we performed an integrated multi-omic analysis, combining transcriptomics and metabolomics. We analyzed global changes in the transcriptome of *xpf-1* worms in mid-life (day 7 of adulthood), when they start displaying increased stress sensitivity but prior to any evidence of increased mortality. In total, 8471 differential expressed genes (*q < 0*.*05*) were identified in *xpf-1* mutants compared to wild-type (N2); 5104 upregulated genes and 3367 downregulated genes (Fig 1a). Enrichment analysis of differentially expressed genes using WormCat tool^27^, showed an upregulation of genes particularly involved in metabolism, stress response, signaling, and mRNA functions (Fig 1b).

**Figure 1.**
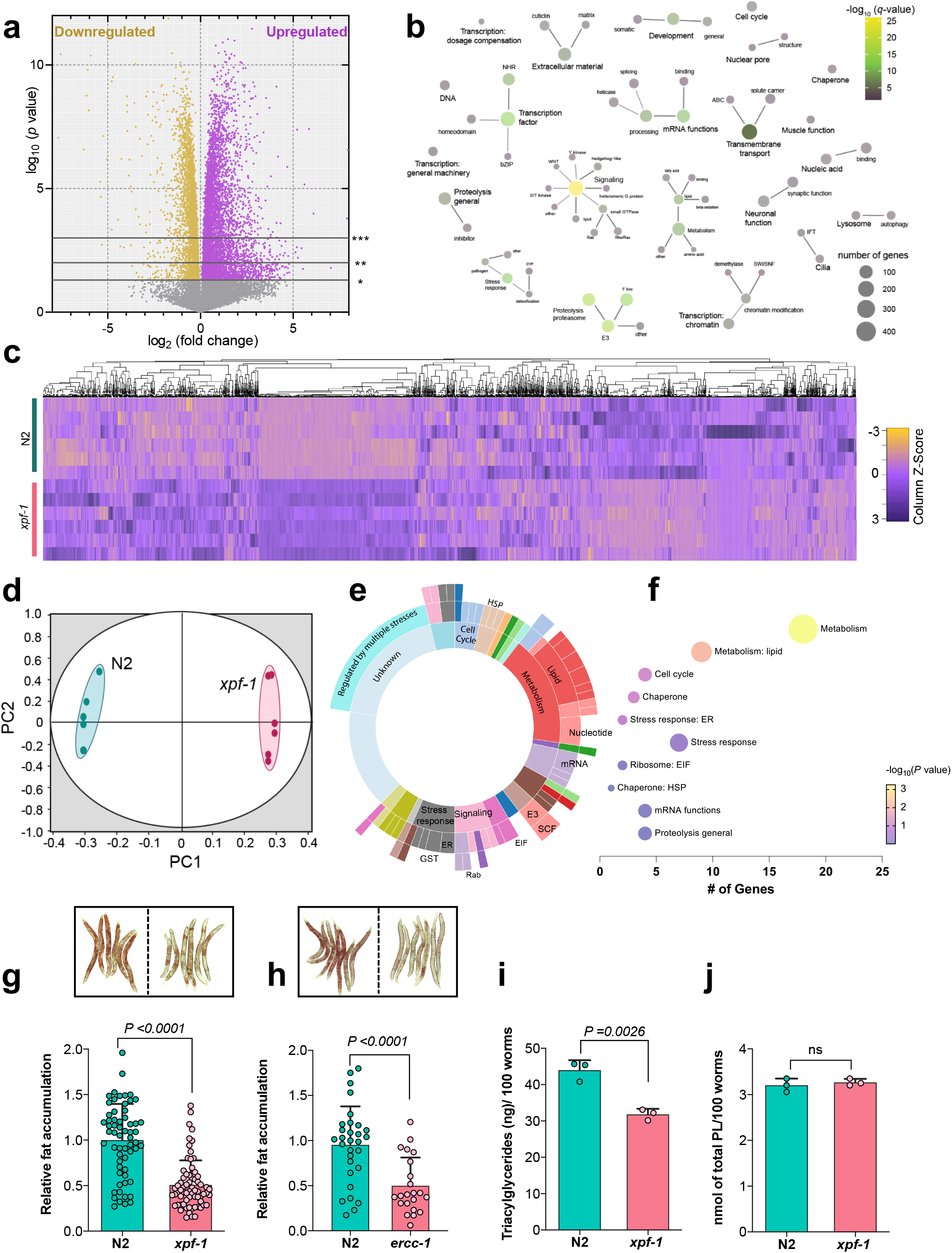
Persistent DNA damage rewires lipid metabolism. **(a)** Volcano plot of differentially expressed genes in *xpf-1 at* day 7 (D7 adulthood). FDR-corrected *p* value calculated using two-tailed Student t test. Horizontal gray lines denote adjusted *p* value cutoffs of 0.05 (*), 0.01 (**), and 0.001(***). The number of genes that are upregulated (purple) or downregulated (gold) significantly (*p* value < 0.05) is indicated. n = 6 biological replicates **(b)** Network diagram showing statistically significant pathways represented by upregulated genes (adjusted p value >0.05) in *xpf-1* at D7. Each node represents the pathway category 1, 2, 3 defined in WormCat. **(c)** Heatmap showing changes in abundance of metabolites in N2, and *xpf-1* **(d)** Dot plot representing the principal component analysis of metabolites in N2 and *xpf-1*. n = 6 biological replicates **(e)** Significantly enriched categories after multi-omic data integration of RNA-seq and metabolomics, performed using Multi-Omics Factor Analysis V2 (MOFA+). **(f)** Bubble plot showing enriched pathways represented by the highly correlated transcriptome and metabolome. The size of the bubble is indicative of the number of genes annotated with WormCat categories; bubbles are color coded according to its FDR corrected *p* values. **(g-h)** Distribution of relative fat depots as measured by ORO staining in **(g)** N2 and *xpf-1* worms. n≥ 61 N2 and *xpf-1* **(h)** N2 and *ercc-1*. n≥ 22 N2 and *ercc-1* **(h)**. Data are Mean ± s.d. Top inset shows representative ORO stained images. 30 worms in N2 Two-way ANOVA, Tukey’s multiple comparison test. **(i)** Total triglycerides (TAG) **(j)** and total phospholipids levels in N2 and *xpf-1*. Data are Mean ± s.d. Two-tailed student t test, n = 3 biological replicates.

Molecular changes in response to stimulants such as DNA damage, begins at the transcriptomic level and are finally translated into metabolomic changes that reflect phenotype. Therefore, we also analyzed the metabolome of N2 and *xpf-1* worms at day 7. Our global metabolomic analysis using high-resolution liquid chromatography mass spectrometry identified 4229 features. We noted that 853 metabolic features were elevated in N2 worms while 1521 were elevated in the *xpf-1* worms (Fig 1c). Multivariate analysis using principal component analysis (PCA) revealed a significant difference between the metabolome of N2 and *xpf-1* worms (Fig 1d). In order to capture the changes in biological processes ensuing with persistent DNA damage at a systemic level we performed a multi-omic data integration of transcriptomics and metabolomics data. Using Multiomics Factor Analysis (MOFA) framework, we first identified transcripts and metabolic features that showed a coordinated increase in *xpf-1* compared to N2^28,29^. Enrichment analysis of highly coordinated sets showed that metabolism, specifically lipid metabolism was significantly upregulated in *xpf-1* mutants (*p*<0.05) (Fig 1e, f). Other pathways identified through enrichment analysis such as cell cycle, mRNA functions, proteolysis were insignificant (*p*>0.05). Together, our multi-omic data suggests that persistent DNA damage rewires metabolism particularly towards lipid metabolism.

Since lipid metabolism was significantly altered in *xpf-1* mutants, we examined temporal changes in fat depots using Oil-red-O (ORO). ORO staining is a semiquantitative method that is routinely used to measure lipid changes in *C. elegans* and it primarily stains neutral lipids such as triglycerides (TAG)^30^. At day 7, *xpf-*1 and *ercc-1* mutants exhibited significantly decreased ORO-staining compared to N2 (Fig 1g, h). Consistent with these results, biochemical quantitation of total TAG levels in *xpf-1* worms showed a substantial decrease (Fig 1i). No changes in the levels of phospholipid (PL) content were observed in these animals (Fig 1j). To determine whether the loss of fat stores was a feature of the ‘*exhaustive*’ phase in response to persistent DNA damage, we analyzed *xpf-1* and *ercc-1* worms as young adults (day 2 of adulthood-‘adaptive’ phase-stress resistant). At day 2, ORO-staining in *xpf-1* was not significantly affected. Consistent with this, we did not observe any changes in TAG and PL levels in young *xpf-1* adults (Supplementary Fig 1 a-c). These findings indicated that lipid stores in DNA-repair mutant worms decline rapidly by mid-life. Although the loss of fat depots was more evident at day 7, the decrease in ORO staining was noticeable in *xpf-1* animals by day 5 (Supplementary Fig 1d). To rule out the possibility of the germline signaling in the regulation of fat depots, we examined ORO staining in *glp-4* mutants, which are defective in germline proliferation^30,31^. Consistent, with *ercc-1* mutants, *ercc-1 glp-4* double mutants showed reduced ORO staining compared to *glp-4* animals, indicating that the loss of fat depots in DNA-repair mutants is not influenced by germline proliferation/germline stem cell signaling (Supplementary Fig 1e). These results suggest that persistent DNA damage results in significant loss of fat stores.

### Lipolysis is augmented upon persistent DNA damage

The exhaustion of lipid stores in DNA-repair mutants can be attributed to either defects in lipid synthesis or an increase in lipid catabolic processes. We first examined lipid synthesis genes by RT-qPCR. FASN-1 encodes a multienzyme complex with six catalytic activities including fatty acid synthase activity, whereas *pod-2* catalyzes the rate-limiting step of generating malonyl-CoA from acetyl-CoA, which is then utilized by FASN-1 to synthesize fatty acids^32^. Both *fasn-1* and *pod-2* were not significantly affected in *xpf-1* mutants (Fig 2a), suggesting that the loss of fat depots observed is likely not due to impaired lipid synthesis but rather an increase in lipid catabolic processes. The majority of the lipid metabolic processes in *C. elegans* occurs in the intestine where the animals store fat as droplet structures mainly composed of TAG^33,34^. During lipid catabolism, first the lipid droplets are hydrolyzed by intestinal lysosomal lipases that breakdown TAGs to free fatty acids (Fig 2b). RT-qPCR analysis showed increased mRNA expression of two lipases, *lipl-1* and *lipl-3*, but no changes in *lipl-4* in *xpf-1* mutants (Fig 2c). Knockdown by RNAi starting at day 1 of adulthood of both *lipl-1* and *lipl-3* restored fat depots in *xpf-1* mutants to that of N2, suggesting that the loss of fat stores observed in *xpf-1* is a result of augmented lipolysis (Fig 2d).

**Figure 2.**
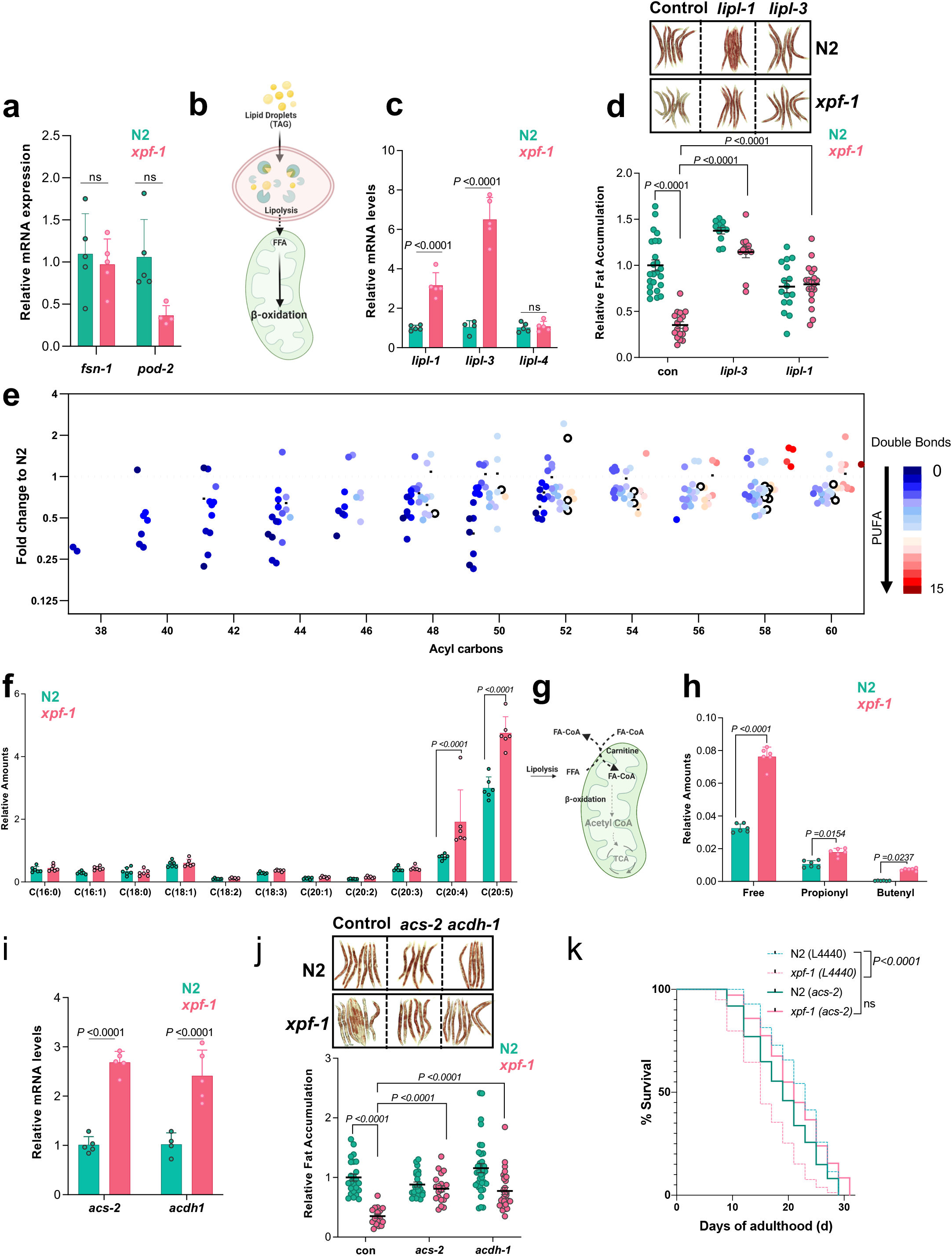
Increased lipolysis and mitochondrial β-oxidation in response to persistent DNA damage. **(a)** Relative mRNA levels of fatty acid synthesis genes in N2 and *xpf-1*. n = 5 biological replicates. Data are Mean ± s.d. Two-way ANOVA with Sidak multiple comparison test. **(b)** Illustration showing triglyceride (TAG) lipid droplets broken down by lipases in lysosomes to release free fatty acids that enter mitochondria for β-oxidation. **(c)** mRNA levels of lysosomal lipases (*lipl*) in N2 and *xpf-1* animals. n = 5, biological replicates. Data are Mean ± s.d. Two-way ANOVA with Sidak multiple comparison test. **(d)** Relative fat stores measured by ORO staining in N2 and *xpf-1* animals fed RNAi against control (L4440), *lipl-3* and *lipl-1*. n>11 worms/condition 2 independent experiments. Data are Mean ± s.d. Two-way ANOVA with Tukey’s multiple comparison test. **(e)** Dot plot showing fold change of TAG lipids in *xpf-1* compared to N2. X axis represents the total acyl carbons in the TAG molecules and color represent the number of double bonds. TAGs with more than 3 double bonds has at least 1 PUFA. Lipids above the dotted line (fold change =1) represent the upregulated lipids and below the dotted line are the downregulated lipids in *xpf-1*. Only significantly changed lipids are shown here (two-tailed Student t test p<0.05). **(f)** Relative amounts of free fatty acids n = 6, biologically independent experiments. Data are Mean ± s.d. two-way ANOVA with Sidak multiple comparison test. **(g)** Schema of free fatty acid mobilization into mitochondria for β-oxidation. **(h)** Levels of free and short-chain carnitine levels in N2 and *xpf-1*. n = 6, biologically independent experiments. Data are Mean ± s.d. two-way ANOVA with Sidak multiple comparison test. **(i)** Relative mRNA levels of β-oxidation genes in N2 and *xpf-1* mutant animals. n = 6, biological replicates. Data are Mean ± s.d. two-way ANOVA with Sidak multiple comparison test. **(j)** ORO staining images (top panel) and quantitation (bottom panel) of relative fat levels in N2 and *xpf-1* fed with RNAi against control (L4440), *acs-2* and *acdh-1*. n >17 worms/condition 2 independent experiments, two-way ANOVA with Tukey’s multiple comparison test **(k)** Lifespan survival of N2 and *xpf-1* grown on control (L4440) or *acs-2* RNAi from day 1 of adulthood. Representative of 3 experiments. Log-rank (Mantel-Cox) test.

Next, to examine the lipid profile changes in response to persistent DNA damage, we performed an in-depth lipidomic analysis. *C. elegans*, like mammals have TAGs that contain mono-unsaturated fatty acids (MUFA), poly-unsaturated fatty acids (PUFA) and saturated fatty acids (SFA). Consistent with an increase in lipolysis, we observed ∼58% of TAG species were significantly reduced in *xpf-1* animals (Fig 2e). Primarily, highly abundant isobaric species of TG 52:2, TG 50:2, TG 54:3, TG 50:3, TG 48:2, TG 50:1, TG 48:1, TG 52:3 were reduced,suggesting hydrolysis of mostly MUFA containing TAGs. Interestingly, ∼13% of TAGs were significantly elevated in *xpf-1* mutants compared to N2. Among the increased TAGs, 78% had at least one PUFA (double bond >3) suggesting a selective synthesis/regeneration of PUFA containing TAGs. Furthermore, lipidomic profiling of free fatty acids in *xpf-1* animals revealed that SFAs, such as palmitic acid (16:0), steric acid (18:0), arachidic acid (20:0) and MUFAs such as, palmitoleic acid (16:1), oleic acid (18:1) and eicosenoic acid (20:1) were unchanged. Major long chain SFAs and MUFAs (≤ 18 carbon) usually enter mitochondria for fatty acid oxidation implying that these free fatty acids may be used for subsequent β-oxidation step (Fig 2g). In contrast, two major PUFAs, arachidonic acid (AA, 20:4) and eicosapentaenoic acid (EPA, 20:5) were elevated in *xpf-1* mutants whereas, minor PUFAs including linoleic acid (18:2), eicosadienoic acid (20:2) and dihomo-γ-linolenic acid (20:3) were unaltered. Taken together, these results indicate an increase in breakdown of lipid depots and enrichment of free-PUFAs in response to persistent DNA damage.

### Increased fatty acid oxidation (β-oxidation) in DNA repair-deficient mutants

Upon lipolysis long chain fatty acids (LCFA) may enter the mitochondria for β-oxidation. However, the mitochondrial membrane is impermeable to LCFA. Hence, LCFA must be conjugated to carnitine to enter the mitochondria. Metabolomic data showed an increase in free, as well as, short-chain acylcarnitine levels (putative identification of butyl and propenyl carnitine) in *xpf-1* mutants suggesting an upregulation in β-oxidation (Fig 2h, Supplementary Fig 2a). Consistent with this metabolomics data, transcriptional levels of genes regulating mitochondrial β-oxidation genes, including *acs-2* and *acdh-1*, were significantly increased in *xpf-1* mutants (Fig 2i). ACS-2 encodes an acyl-CoA synthetase that activates fatty acids for transport into the mitochondria^35,36^, whereas, ACDH-1, encodes a mitochondrial short-chain acyl-CoA dehydrogenase^37^ that catalyzes the initial enzymatic reaction of β-oxidation. RNAi mediated knockdown of both *acs-2* and *acdh-1* in *xpf-1* adults rescued the loss-of-fat phenotype (Fig 2j), suggesting that the fat stores are depleted in *xpf-1* mutants as a consequence of increased β-oxidation.

We next performed a RNAi screen for potential transcription factors that could be regulating lipid metabolism in *xpf-1* mutants. Interestingly, none of the tested transcriptional regulators involved in lipid metabolism, including *hlh-30* (mammalian homologue-TFEB), *nhr-49* (PPARα), *nhr-80* (HNF4), *sbp-1* (SREBP) and *mdt-15* (MED15), rescued the DNA damage-induced loss of fat depots (Supplementary Fig 2b). Overall, these results indicate that persistent DNA damage drives a metabolic switch to β-oxidation, independent of the canonical lipid metabolic transcription factors. We then examined whether the induction of mitochondrial β-oxidation in DNA repair-deficient mutants is associated with their shortened lifespan. RNAi knockdown of *acs-2* not only restored fat accumulation, but also recued the shortened lifespan in *xpf-1* animals (Fig 2k). Together, these data suggested that there is an increase in β-oxidation in *xpf-1* mutants and inhibiting mitochondrial β-oxidation rescues lifespan.

### Persistent DNA damage increases β-oxidation for generation of acetyl-CoA

β-oxidation breaks down fatty acids to generate acetyl-CoA, a metabolite that plays a pivotal role not only in energy production and macromolecular synthesis, but also acts as a sole donor for acetylation^17^. Acetyl-CoA can have multiple fates including entering the TCA cycle to ultimately generate ATP. Metabolomic profiling of TCA metabolites showed that the levels of citrate, fumarate and malate were not significantly affected in *xpf-1* mutants (Fig 3a). Consistent with these results, the mRNA levels of genes involved in the production of these metabolites were not affected (Fig 3b). Key glycolytic metabolites such as hexose, hexose-6-phosphate and pyruvate, which link to acetyl-CoA production and the TCA cycle, were also not altered (Fig 3a). Furthermore, amino acid levels were not changed between N2 and *xpf-1* animals, except for valine and glutamine (Supplementary Fig 3a). Both valine and glutamine can feed into TCA cycle to increase the levels of succinate^38,39^. Succinate, a key metabolite that connects TCA cycle to electron transport chain, was elevated in *xpf-1* mutants (Fig 3a) and was paralleled with decrease in succinate dehydrogenase B and D enzymes (Fig 3b). It must be noted that the levels of metabolites downstream of succinate, such as malate and fumarate were unaltered in *xpf-1* mutants, suggesting that the increase in valine and glutamine has little effect on the overall output of the TCA cycle. These data suggest that the steady state TCA cycle is not altered markedly upon persistent DNA damage.

**Figure 3.**
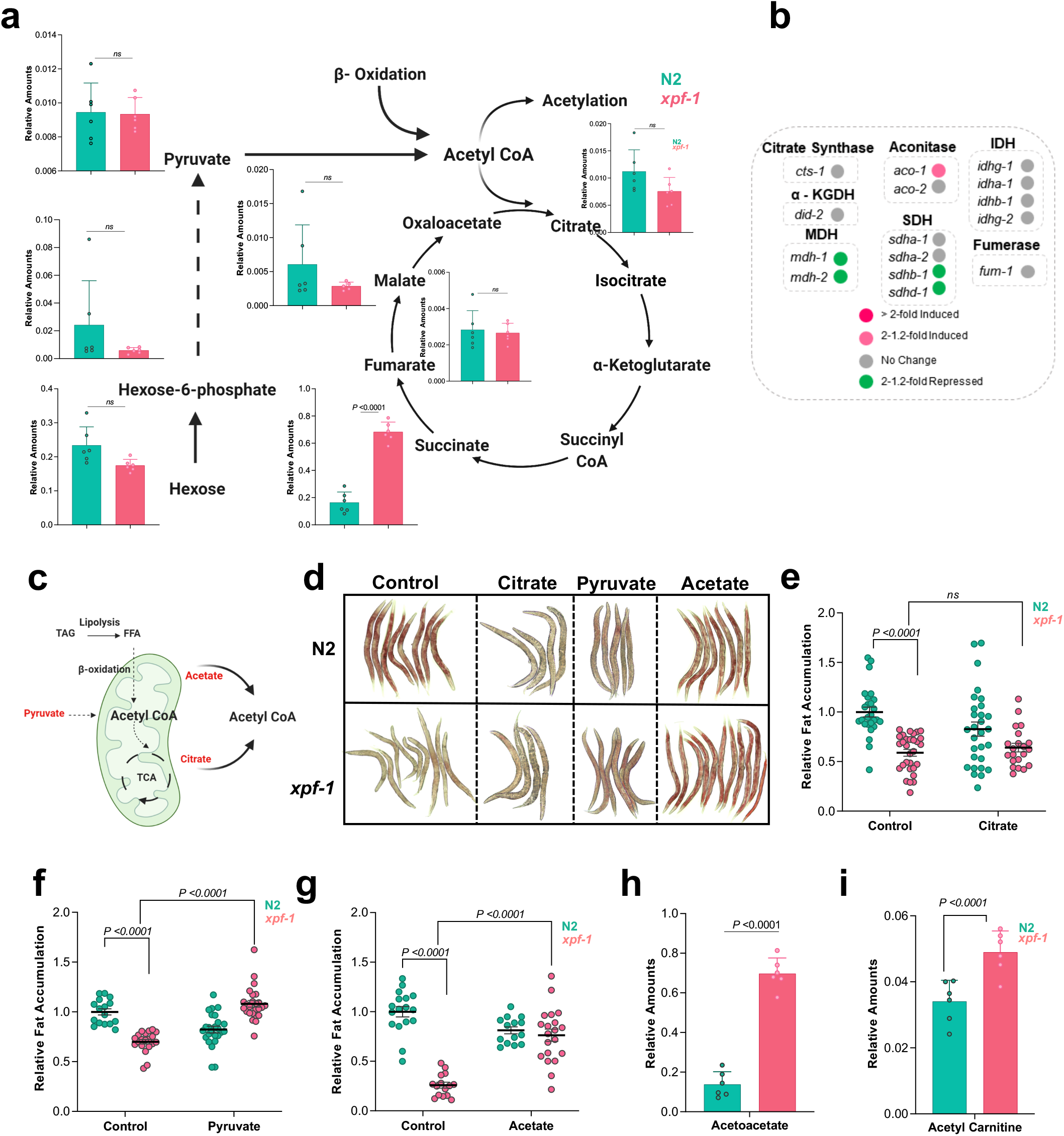
Increased β-oxidation in *xpf-1* is used to generate acetyl-CoA. **(a)** Glycolysis and Tricarboxylic acid cycle (TCA) metabolite levels in N2 and *xpf-1*. n = 6, biologically independent experiments, Student’s two-tailed t test. **(b)** TCA metabolic gene expression profiles between N2 and *xpf-1*. **(c)** Schema illustrating sources of acetyl-CoA production. Metabolites in red when supplemented exogenously can generate acetyl-CoA. **(d)** Representative ORO stained images and **(e-g)** quantitation of fat stores in N2 and *xpf-1* in control or supplemented with **(e)** citrate, **(f)** pyruvate and **(g)** acetate. n> 14 worms/condition 2-4 independent experiments, Data are Mean ± s.d. two-way ANOVA with Tukey’s multiple comparison **(h)** Levels of acetoacetate, a product of acetyl-coA in N2 and *xpf-1*. n = 6, biologically independent experiments, Student’s two-tailed t test. **(i)** Relative amount of acetyl carnitines in N2 and *xpf-1* worms. n = 6 biologically independent experiments. Data are Mean ± s.d. Student’s t test.

Next, we wanted to directly test whether the increase in β-oxidation leading to loss of fat stores in *xpf-1* mutants was primarily for the generation of acetyl-CoA. We reasoned that if *xpf-1* mutants were upregulating β-oxidation to generate acetyl-CoA, then supplementation with metabolites known to directly produce acetyl-CoA, would rescue β-oxidation and thus subsequent loss of fat stores. We supplemented the N2 and *xpf-1* nematodes with three metabolites known to directly produce acetyl-CoA: citrate, pyruvate and acetate (Fig 3c). Citrate which can be converted into acetyl-CoA either by ATP-citrate lyase (*acly*/ACLY) or through the TCA cycle did not rescue the fat stores in *xpf-1* mutant suggesting both these routes are unable to generate acetyl-CoA (Fig 3d, e). Pyruvate and acetate are converted to acetyl-CoA by pyruvate dehydrogenase (*pdh*/PDH) and acyl-coenzyme A synthetase (*acs-19*/ACSS2), respectively^40,41^. Supplementation with either pyruvate or acetate rescued the loss of fat stores in *xpf-1* mutants (Fig 3d, f, g). Interestingly, we did not observe a substantial difference in the total acetyl-CoA concentration between *xpf-1* and N2 worms (Supplementary Fig 3b, c). However, we observed an increase in acetyl-CoA dependent metabolites. A number of acetylated amino acids and amino acid derivatives were increased in *xpf-1* worms (Supplementary Fig 3d, f-n). Furthermore, we observed an increase in acetoacetate (putative identification) in *xpf-1* worms through metabolomic fragmentation analysis (Fig 3h) which is a product of mitochondrial compartmentalized acetyl-CoA via ketogenesis. Similarly, we observed an increase in acetyl-carnitine levels in the *xpf-1* mutants, suggesting the potential export of acetyl-CoA from the mitochondria (Fig 3i). Taken together, our findings suggest that DNA repair mutants drive mitochondrial β-oxidation in order to generate acetyl-CoA.

### β-oxidation derived acetyl-CoA promotes histone hyperacetylation

Acetyl-CoA is a sole donor of acetyl groups for histone and non-histone acetylation^40,42,43^ (Fig 4a). Previously, it was reported that mitochondrial β-oxidation is the predominant contributor for histone acetylation, chromatin regulation and gene transcription^43,44^. Acetyl-CoA is impermeant to the inner mitochondrial membrane and needs to be transported into or generated locally in the nucleus for histone acetylation^45^. Acetyl-carnitines are known to shuttle acetyl-CoA from mitochondria into the nucleus^46^. We next examined two key histone acetylation sites associated with DNA damage, as well as aging. Histone 4 lysine 16 acetylation (H4K16ac) opens chromatin by reducing the inter-nucleosomal interactions^47^. On the other hand, histone 3 lysine 9 acetylation (H3K9) acetylation is elevated in the region surrounding the transcription start sites of active genes indicating its role in transcriptional initiation^48,49^ (Fig 4a,b). H4K8/16ac and H3K9ac marks were significantly increased in *xpf-1* mutants compared to N2 (Fig 4b). Previously, it was shown that variations in H3/H4 acetylation levels positively correlated with nuclear size^50^. Since we observed H4K8/16 hyperacetylation, which leads to open chromatin conformation, we examined the nuclear size in intestinal cells of *xpf-1* nematodes. DAPI stained intestinal nuclei in *xpf-1* mutants were enlarged by ∼40% (N2:169.3 ± 8.0 µm^2^, *xpf-1*: 237.6 ± 9.8 µm^2^), while the number and DNA content of the intestinal nuclei remained unchanged, thus indicating a ‘loose’ chromatin structure (Fig 4c, f). Interestingly, knockdown of *acs-2* in these animals reduced H4K8/16 hyperacetylation and restored nuclear size towards control levels (189.1±10 µm^2^; Fig 4d-f). These findings suggest that the upregulated β-oxidation in *xpf-1* mutants drives hyperacetylation of histones affecting global changes in chromatin structure. Increase in histone acetylation are typically associated with enhanced gene expression. Consistent with this, our RNA-seq data showed an increase in global gene expression levels (Fig 1a). Specifically, an upregulation of ‘immune-like signature genes’ including C-type lectins (*clecs*) and cytochrome P450s (*cyp*) genes was observed in *xpf-1* worms (Supplementary Fig 4a). CYPs in particular are stress response genes that belong to a family of diverse monooxygenases, which oxidize fatty acids, sterols and endogenous compounds. Upregulated CYP expression in *xpf-1* mutants was substantiated with RT-qPCR data (Fig 4g). Thus, persistent DNA damage promotes histone hyperacetylation and upregulation of an immune-effector gene transcription program.

**Figure 4.**
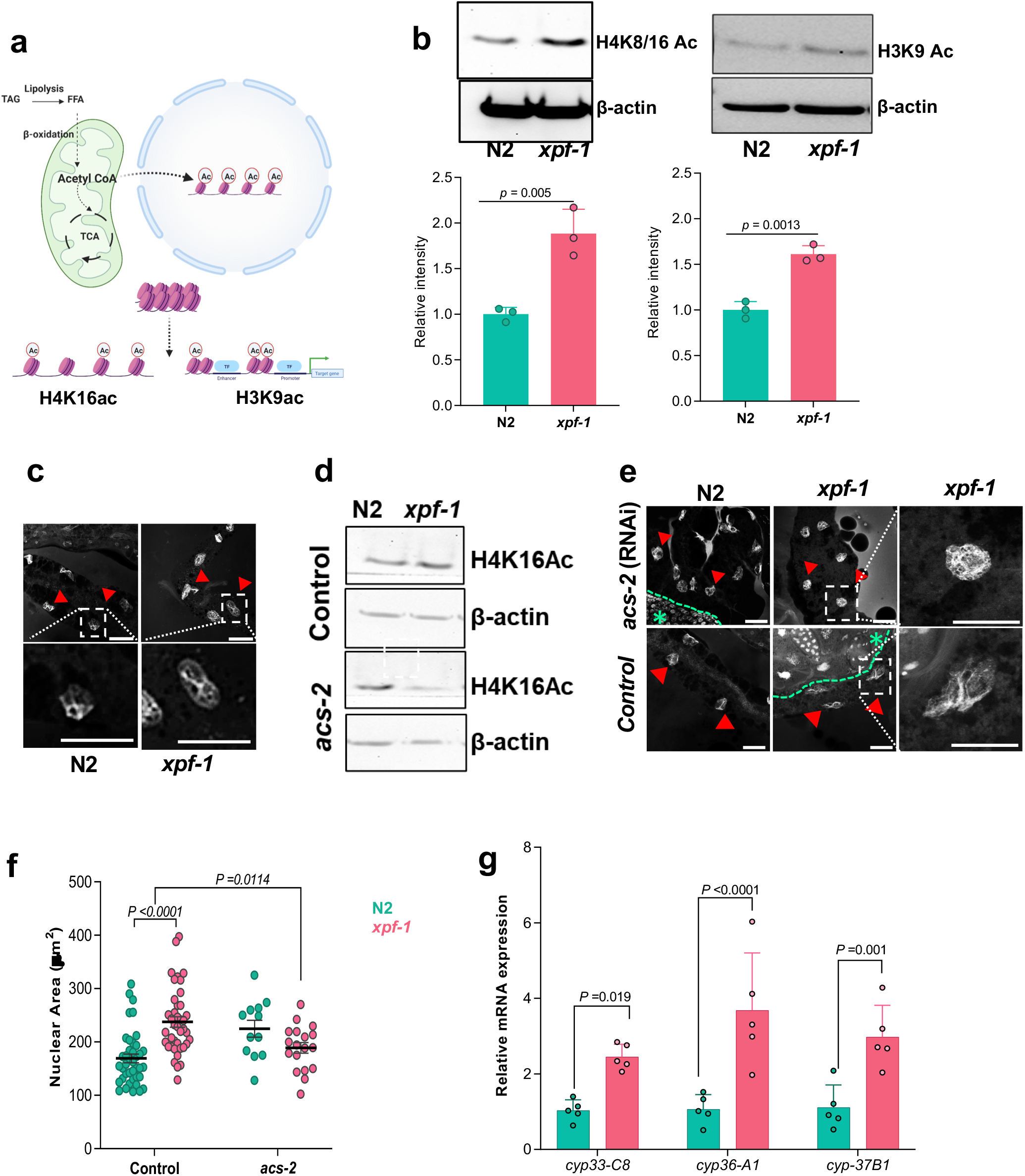
Increased β-oxidation drives histone acetylation upon persistent DNA damage. **(a)** Illustration showing increased β-oxidation leads to an increase in acetyl-CoA pools that is used for histone acetylation in the nucleus. H4K16ac opens up chromatin structure and H3K9ac marks are present at active gene promoters. **(b)** Immunoblots of histone H4K8/16Ac and H3K9Ac in N2 and *xpf-1* with anti-β-actin as loading control (top panel). Quantitation of histone H4K8/16 and H3K9 levels relative to β-actin (bottom panel). n = 3. Data are Mean ± s.d. two-tailed Student’s *t* test **(c)** Representative images of DAPI staining of intestinal nuclei in N2 and *xpf-1* mutant animals fed with L4440 (control) **(d)** Immunoblots of histone H4K8/16Ac in N2 and *xpf-1* treated with (L4440 RNAi) or (*acs-2* RNAi) with anti-β-actin as loading control. **(e)** Representative images of DAPI staining of intestinal nuclei in N2 and *xpf-1* mutant animals fed with L4440 (control) or *acs-2* RNAi. Dash rectangles highlight the areas enlarged and shown below. Scale bars= 25 μm. The red arrow indicates the intestinal nucleus; the area with green dotted line or asterisk denotes nucleus of germ cells. **(f)** Quantitation of intestinal nuclear cross-sectional area in N2 and *xpf-1* mutant animals grown on L4440 (control) and *acs-2* RNAi. n =40 for N2 and *xpf1*(control), 12 for N2(*acs-2*) and 18 for *xpf-1*(*acs-2*). Data are Mean ± s.d. two-way ANOVA, Tukey’s multiple comparison test. **(g)** Relative mRNA levels of CYP genes in N2 and *xpf-1*. n = 5 biological replicates. Data are Mean ± s.d. Two-way ANOVA with Sidak multiple comparison test.

### MYS-1 controls histone hyperacetylation in response to persistent DNA damage

Histone acetylation is a highly regulated process governed by two enzyme groups, histone acetyl transferases (HAT) and histone deacetylases (HDAC)^51^. HATs transfer acetyl groups from acetyl-CoA, to an ε-amino group of a histone lysine residues^52^. On the other hand, HDACs remove acetyl groups from acetylated histones^53^. Balance between HATs and HDACs play a crucial role to regulate chromatin dynamics, gene expression and to maintain homeostasis^54^ (Fig 5a). To determine the HATs involved in the hyperacetylation of histones in *xpf-1* mutants, we performed an RNAi screen targeting putative HATs. We reasoned that since acetyl-CoA is an obligatory cofactor of HATs, knockdown of the involved HAT would avert the increased demand for acetyl-CoA. This would thus relieve the need for augmented β-oxidation and rescue the loss of fat stores in *xpf-1* nematodes. We knocked down three subfamilies of HAT-1 class previously known to play a role in DNA damage recognition and repair: *pcaf-1* (mammalian homologue PCAF), *mys-1* (Tip60), *cbp-1* (CBP/p300)^55^. Only *mys-1* knockdown resulted in the rescue of fat stores in *xpf-1* mutants suggesting its role in acetylation, using β-oxidation-derived acetyl-CoA (Fig 5b). Consistent with these findings, the level of H4K8/16ac was also reduced upon *mys-1* knockdown (Fig 5c). We next examined whether the increased nuclear size in *xpf-1* mutants was also rescued upon suppression of *mys-1*. Indeed, DAPI staining of intestinal nuclei in *xpf-1* (*mys-1* RNAi) exhibited compact nuclei (Fig 5d, e). Since *mys-1* knockdown restored fat depots, decreased histone hyperacetylation and nuclear size, we next examined its effects on lifespan of *xpf-1* mutants. Suppression of *mys-1* rescued the shortened lifespan of *xpf-1* worms (Fig 5f). Next, we assessed the HDACs involved in the DNA damage driven metabolic-epigenetic axis. Since HDACs deacetylate histones, inhibition of involved HDAC would prevent additional demand for acetyl-CoA, and therefore rescue fat stores. A targeted RNAi screen for histone deacetylases: *hda-1, hda-3* and *sir-2*.*1* revealed only *sir-2*.*1* led to increased fat stores in *xpf-1* mutants (Fig 5g, h). As expected, *sir-2*.*1* knockdown did not affect the intestinal nuclei size in *xpf-1* animals (Fig 5i, j). Likewise, *sir-2*.*1* knockdown did not alter the shortened lifespan of *xpf-1* mutants (Fig 5k). Overall, these results indicate that an increase in β-oxidation is tightly coupled with histone hyperacetylation, and this drives aging in response to persistent DNA damage.

**Figure 5.**
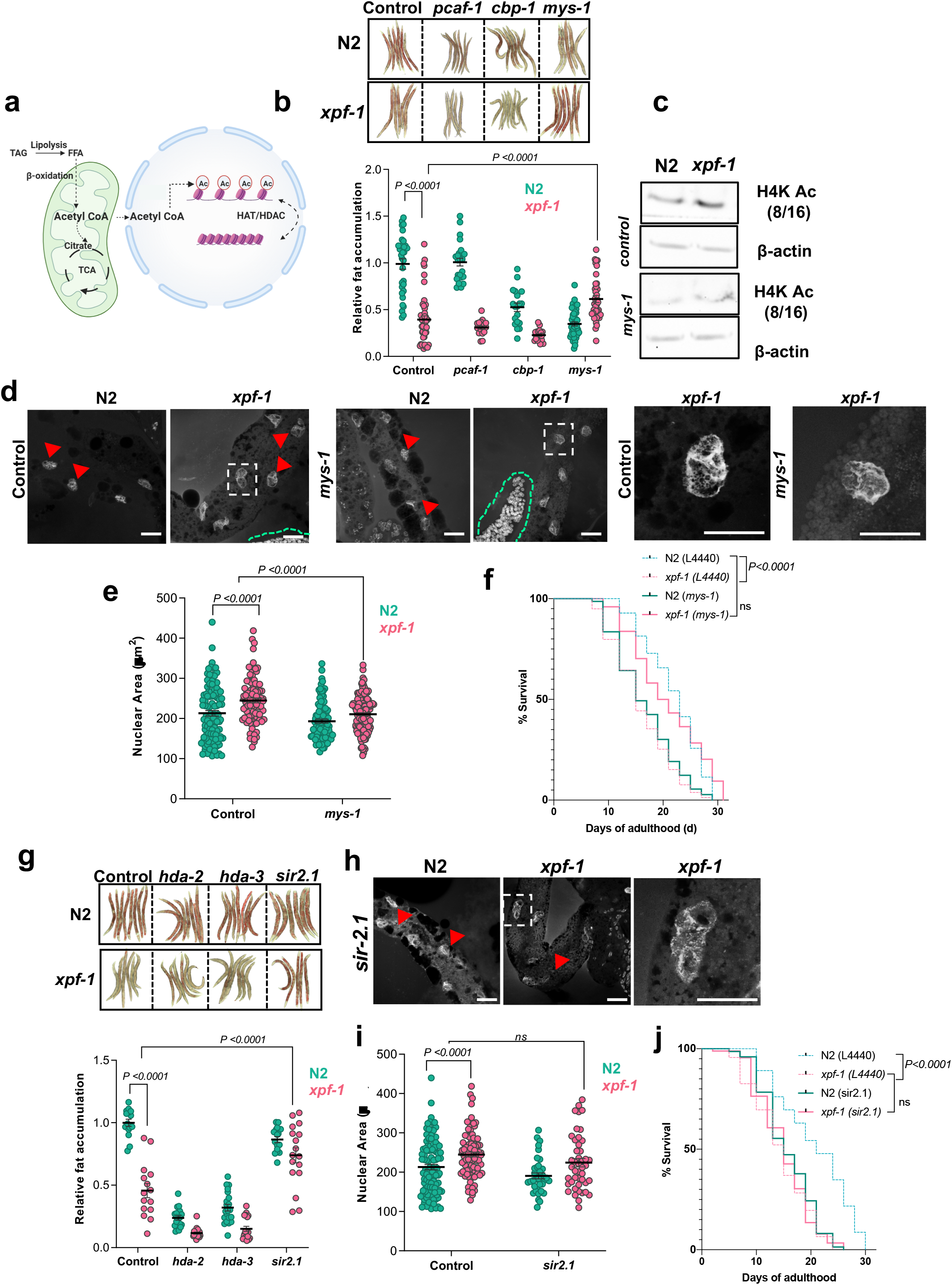
MYS-1 and Sir2.1 control histone hyperacetylation in response to persistent DNA damage. **(a)** Illustration showing acetylation, deacetylation of histones by histone acetyl transferases (HATs) and histone deacetylase (HDACs), respectively **(b)** Representative images and quantitation of fat depots by ORO staining in N2 and *xpf-1* animals fed with control (L4440) and *pcaf-1, cbp-1* or *mys-1* HAT RNAi. Data are Mean ± s.d. two-way ANOVA, Tukey’s multiple comparison test. **(c)** Immunoblots of H4K8/16Ac and H3K9Ac in N2 and *xpf-1* animals fed with L4440 and *mys-1* RNAi. **(d)** Representative image of DAPI staining of intestinal nuclear cross sectional area in N2 and *xpf-1* with control (L4440) and *mys-1* RNAi and **(e)** quantitation. Data are Mean ± s.d. two way ANOVA, Tukey’s multiple comparison test. **(f)** Lifespan survival of N2 and *xpf-1* with control (L4440) and *mys-1* RNAi. Log-rank (Mantel-Cox) test. **(g)** Representative images and quantitation of lipid stores by ORO staining in N2 and *xpf-1* with control (L4440) and *hda-2, hda-3* and *sir2*.*1* HDAC RNAi. Data are Mean ± s.d. two-way ANOVA, Tukey’s multiple comparison test. **(h)** Representative image of DAPI staining of intestinal nuclear cross-sectional area in N2 and *xpf-1* fed with *sir-2*.*1* RNAi and **(i)** quantitation. Data are Mean ± s.d. two-way ANOVA, Tukey’s multiple comparison test. **(j)** Lifespan survival of N2 and *xpf-1* with control (L4440) and *sir-2*.*1* RNAi. Log-rank (Mantel-Cox) test.

### Lipidome changes in *xpf-1* worms

Thus far our data suggests that persistent DNA damage rewires lipid metabolism to promote histone hyperacetylation through MYS-1. Furthermore, we observed transcriptional upregulation of CYP genes, and an accumulation of PUFAs. CYPs can act on PUFAs and generate bioactive lipids^56^. Therefore, we wanted to closely examine the lipidome alterations to identify bioactive lipid species in *xpf-1* mutants. We performed untargeted lipidomic profiling of N2, *xpf-1* worms treated with and without *mys-1* RNAi. Using a reversed-phase high-resolution liquid chromatography tandem mass spectrometry (HR-LC-MS/MS) we identified 12 lipid classes. Negative ion mode was used to identify most of the phospholipids including oxidized phospholipids, as well as free fatty acids and oxidized free fatty acids (Fig 6a), whereas positive ion mode was used to quantify the triacylglycerol (TAG) species. In total, we identified 793 lipid species including 354 TAG species and 175 oxidized free fatty acid species. Among phospholipids, 57 phosphatidyl ethanolamines, 52 cardiolipins, 47 phosphatidylcholines and 23 lysophospholipids were identified (Fig 6b). As expected, knockdown of *mys-1* in *xpf-1* mutants restored TAG levels (Fig 6c, Supplementary Fig 6a). Similarly, PUFAs such as arachidonic acid (AA) and eicosapentaenoic acid (EPA) that were elevated in *xpf-1*, were substantially reduced after *mys-1* knockdown (Fig 6d). Since PUFAs are susceptible to lipid peroxidation, we next specifically examined for oxidized free fatty acids. At least 61% oxidized fatty acid species were significantly elevated in *xpf-1* mutants suggesting an increase in lipid oxidation in *xpf-1* mutants (Fig 6e *top panel*). However, upon *mys-1* knockdown, 57% oxidized fatty acids were significantly decreased in *xpf-1* mutants (Fig 6e bottom panel). These oxidized fatty acids were initially identified based on the exact mass, retention time and its ability to lose CO2 during fragmentation analysis. Subsequent fragmentation analysis and comparison with standards revealed the three prominent lipid mediators, prostaglandin E2, 11,12-EpTrE (11, 12-epoxy-eicosatrienoic acid) and leukotriene B4, were elevated in *xpf-1* mutants (Fig 6f). *C. elegans* lack lipid-oxidizing enzymes such as lipoxygenases and cyclooxygenases, however, several oxidized free fatty acids including prostaglandins have been reported in these models. Studies suggest CYP genes are involved in the synthesis of oxidized fatty acids in worms (Fig 6g). Thus, we hypothesized that the upregulation of CYP genes in *xpf-1* mutants is regulated by *mys-1* mediated histone acetylation and gene transcription. Indeed, knocking down *mys-1* decreased *clec* and *cyp* gene expression, and correlated with the reduction in lipid mediators (Figure 6h, Supplementary Fig 4c).

**Figure 6.**
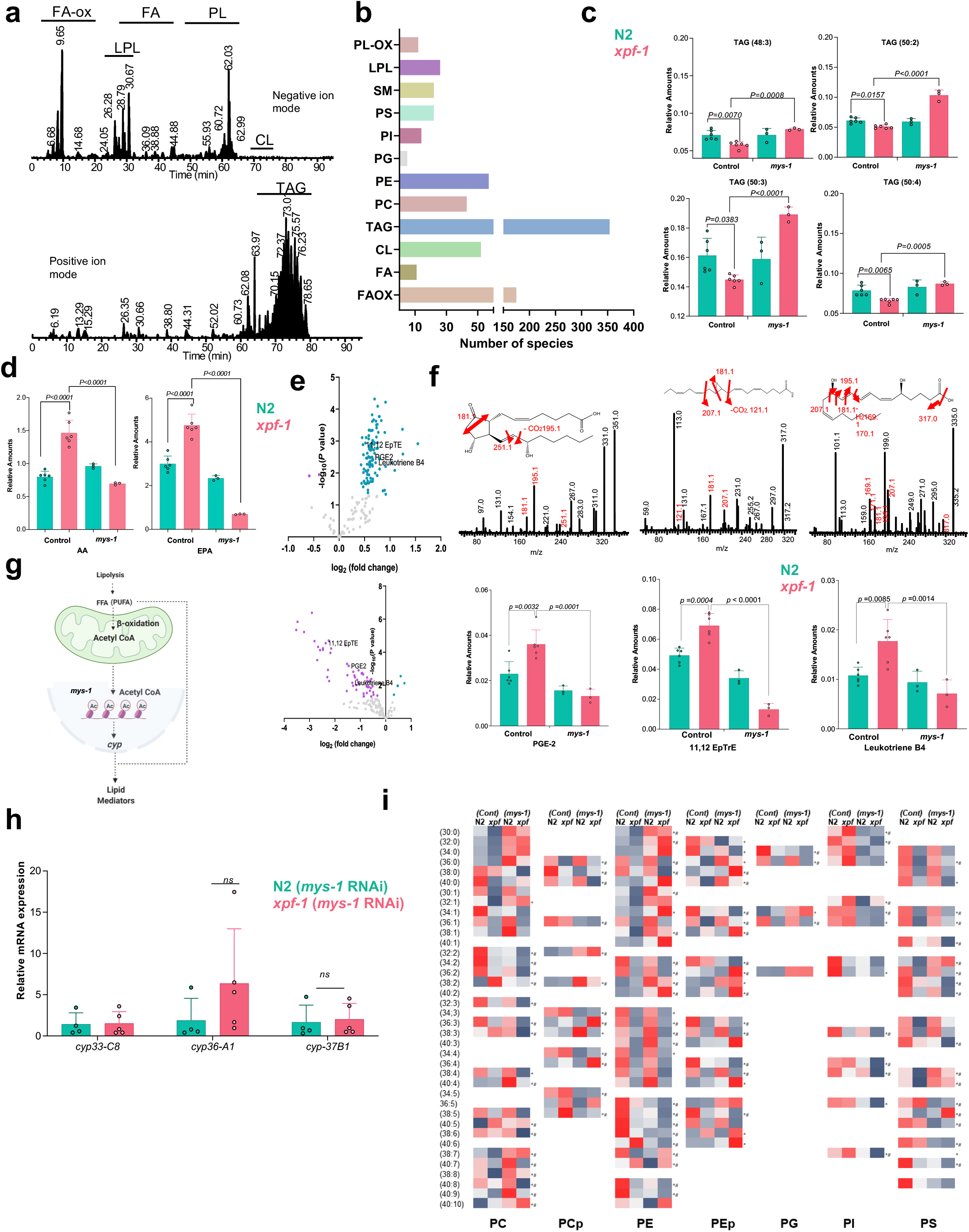
Persistent DNA damage increases lipid mediators. **(a)** Extracted ion chromatography from high resolution liquid chromatography mass spectrometry of *C. elegans* in negative ion mode (*top* panel) and positive ion mode (*bottom* panel). Lipid classes such as oxidized free fatty acid, fatty acids, lysophospholipids (LPL), phospholipids (PL) including (CL) was quantified from negative ion mode, whereas triacylglycerol was quantified from negative ion mode. **(b)** Bar graph showing number of species identified in each lipid categories. **(c)** Bar graphs showing levels of major triacylglycerol species that were significantly reduced in *xpf-1* mutants and restored upon knocking down *mys-1*. Data are Mean ± s.d.. two-way ANOVA, Tukey’s multiple comparison test. **(d)** Bar graph showing levels of arachidonic acid (AA) and eicosapentaenoic acid (EPA) N2, *xpf-1* worms with and without *mys-1* RNAi. Data are Mean ± s.d. two-way ANOVA, Tukey’s multiple comparison test. **(e)** Volcano plot showing the differences in oxidized free fatty acid levels in *xpf-1* vs N2 worms (top panel) and *xpf-1*(*mys-1 RNAi*) vs *xpf-1* worms. **(f)** Distribution of three lipid mediators prostoglandin-E2, 11-12 EpTrE, and leukotriene B4, identified in the *C. elegans* lipidome. Top panel shows the fragmentation spectrum of each lipid mediator, and the bottom panel shows its distribution. Data are Mean ± s.d. two-way ANOVA, Tukey’s multiple comparison test. **(g)** Working model showing excess β-oxidation drives histone hyperacetylation and production of lipid mediators in *xpf-1* **(h)** Levels of *cyp* genes in N2(*mys-1 RNAi*) and *xpf-1*(*mys-1 RNAi*) worms measure by qRT-PCR. Data are Mean ± s.d. two-way ANOVA, Tukey’s multiple comparison test. **(i)** Heat map showing the distribution of phospholipids in N2, *xpf-1*, N2(*mys-1*), *xpf-1*(*mys-1*) worms. * p< 0.05, *xpf-1* vs N2 ^#^ *p*< 0.05, *xpf-1*(*mys-1*) vs *xpf-1*. Student’s t test n = 6 for N2, *xpf-1*; 3 for N2(*mys-1*) and *xpf-1*(*mys-1*)

Phospholipids are the major sources for PUFAs in biological system. Since we observed significant changes in free PUFA and oxidized fatty acids levels, we then analyzed the phospholipidome of *xpf-1* mutant. The majority of PUFA containing phosphatidylcholines and phosphatidylethanolamines were significantly decreased in *xpf-1* mutants. Additionally, the levels of all identified lysophosphotidyl ethanolamine species were decreased in *xpf-1* mutants. On the other hand, levels of phosphatidylinositol were increased in *xpf-1* mutants, suggesting alterations in phospholipid metabolism in these mutants. Knockdown of *mys-1* in *xpf-1* mutants significantly increased several prominent SFA and MUFA containing PE species such as 30:0, 32:0, 30:1, 38:1, 38:2, whereas levels of predominant PUFA containing PE species such as 36:5, 38:5, 40:5, 38:6, 40:6 were significantly reduced (Figure 6i). Taken together these results suggest that suppression of *mys-1* during persistent DNA damage facilitates a reduction in oxidizable lipid substrates. Sphingomyelins are another category of lipids that showed differences in *xpf-1* after *mys-1* treatment. All sphingomyelin species were elevated in *xpf-1* mutant while *mys-1* treatment decreased all, but one species (SM 32:0) (Supplementary Fig 6b). Overall, these findings revealed that persistent DNA damage caused a global shift in lipid metabolism with the most significant alterations occurring in TAGs and free PUFAs.

## DISCUSSION

In this study, we describe a mechanism through which persistent DNA damage alters the metabolic-epigenetic axis and promotes an immune-like response. We show that persistent DNA damage reprograms metabolism to mitochondrial β-oxidation for generating acetyl-CoA. This acetyl-CoA promotes aberrant histone hyperacetylation, chromatin opening and subsequent expression of immune-effector genes. Furthermore, our data indicates that persistent DNA damage-driven metabolic rewiring increases free PUFAs that can be metabolized by the CYP gene family resulting in pro-inflammatory lipid mediators. The HAT, *mys-1*, is responsible for the persistent DNA damage-driven epigenome alterations and age-related decline.

Fat storage and breakdown plays an important role in maintaining health. In humans, excessive fat depots are usually associated with chronic disease. However, several studies also point to the fact that loss of fat stores is an important risk factor in older adults. It is associated with frailty, higher hospital admissions, functional decline and poor quality of life^57-59^. Indeed, increase in fat storage has also been associated with longevity in both invertebrates, as well as mammals^60^. Interestingly, several animal models with persistent DNA damage have loss of fat depots, consistent with our results^18,19,61^. Here, we provide mechanistic insights into how depleted fat depots are a byproduct of excessive β-oxidation and inhibition of this process is beneficial in the context of persistent DNA damage.

Emerging evidence shows that nuclear DNA damage and metabolism are inextricably linked^62^. Acute genotoxic stress tightly and transiently coordinates metabolic events to promote repair and maintain homeostasis^12^. However, the cellular and metabolic events in response to persistent DNA damage are only recently being explored. Persistent DNA damage in the liver and hepatocytes of *Ercc1*^*-/*^Δ mice was shown to inhibit glycolysis and favored NADPH production through the pentose phosphate shunt^63^. A recent study also showed that in response to both acute and persistent genotoxic stress, an increase in mitochondrial β-oxidation feed into oxidative phosphorylation in a cell-autonomous manner^19^. In this study, inhibition of β-oxidation through the CPT inhibitor, etomoxir, seemed to have detrimental effects. However, it is important to note that in this case, etomoxir treatment was performed after an acute exposure to a DNA damaging agent. We have, in fact, reproduced these results and believe that with acute DNA damage, a transient shift towards β-oxidation is beneficial and required for DNA repair (data not shown). In contrast, during persistent DNA damage, our study shows that genetic inhibition of mitochondrial β-oxidation in DNA repair-deficient animals restored fat depots, suppressed histone hyperacetylation and extended lifespan. Therefore, in the context of persistent DNA damage that occurs with age, maintaining β-oxidation could be detrimental.

Several studies show that specific inhibition of mitochondrial β-oxidation, significantly reduces acetyl-CoA levels, while basal mitochondrial respiration and ATP remain unaffected^64,65^. Moreover, a recent study showed that lipid derived acetyl-CoA is mostly responsible for histone acetylation, chromatin regulation and gene transcription^46^. Consistent with this, we observed that the increased mitochondrial β-oxidation was intimately linked with histone hyperacetylation. Emerging evidence supports a role for histone acetylation in aging. For example, H4K16ac accumulates during yeast replicative aging^66^ and leads to a loss of H4 histone in proliferating cells. Although we did observe an increase in H4K16ac, this was not associated with a general loss of histones in DNA repair-deficient mutant *C. elegans*. H4K16ac has also been observed in aging brains, but a decrease in this histone mark was associated with an age—related pathology, Alzheimer’s disease (AD), suggesting that it is critical to have an in-depth understanding of both global and local changes of these histone modifications. Future studies to discern the patterns of these histone acetylation changes need to be performed to understand how they contribute to age-related decline. Interestingly, histone hyperacetylation has been previously shown to increase reactive oxygen species (ROS)^67^, whereas, deacetylation of histone H3 led to upregulation of autophagy genes and longevity^68,69^. Consistent with this, overexpression of the sirtuin family of histone deacetylases have long been linked to longevity^70^. These studies suggest that histone hyperacetylation may have a causal role in promoting biological aging. Our study now provides a link between histone hyperacetylation and an increase in chronic immune-like response.

Since, persistent DNA damage drives rewiring of lipid metabolism, we comprehensively analyzed lipid alterations in this context. The levels of two related lipid classes, free polyunsaturated fatty acids (PUFAs) and oxidized free fatty acids correlate with the lifespan of *C. elegans*. Both of these lipids were elevated in the short-lived *xpf-1* mutants, but reduced upon knockdown of *mys-1*, suggesting that the HAT, MYS-1, is the regulator of the metabolic-epigenetic axis in response to persistent DNA damage. A general consensus is that the higher ratio of MUFA-to-PUFA is crucial in maintaining healthy lifespan^71,72^. Supplementation, as well as genetic modification of PUFA producing enzymes support this notion^60,73-75^. In our study, the MUFA/PUFA ratio in the free fatty acid pools decreased dramatically, while changes in the esterified forms was negligible. This suggests that free fatty acids are likely more capable of regulating lifespan than the esterified forms. Among PUFAs, generally ω-3 PUFA such as ALA and EPA are beneficial^75,76^ while, ω-6 PUFA are often detrimental. The majority of the oxidized free fatty acids derived from ω-6 PUFAs are pro-inflammatory lipid mediators, which leads to a decline in cell function or to cell death^77,78^. In contrast, the majority of lipid mediators derived from ω-3 fatty acids signal anti-inflammatory events and are pro-survival signals^79^.

In DNA repair-deficient mutants we observed an increase in CYP genes. While *cyp* genes are usually involved in xenobiotic and endobiotic response, in the presence of free PUFAs, they can exhibit dichotomous roles due to their ability to produce pro-inflammatory lipid mediators^80^. With persistent DNA damage we observed elevated levels of PUFAs suggesting that the increased oxidized fatty acids are through the enzymatic action of CYPs on PUFA. Interestingly, although both ω-3 and ω-6 PUFAs are elevated in *xpf-1* mutants, ∼95% of the elevated lipid mediators are derived from the ω-6 PUFA, AA. Taken together, the results indicate a strong pro-inflammatory condition in *xpf-1* mutants. Supporting this, the transcriptomic analysis also showed elevation of several immune response-like genes possibly leading to a scenario of chronic inflammation. There is accumulating evidence that suggests that persistent DNA damage leads to increased inflammation, and blunting this immune response is beneficial for health. Our multi-omic study has now explored the inter-dependence of the metabolic, epigenomic and inflammatory changes. Our data suggests that the shift in metabolism to β-oxidation, which leads to loss of fat depots, is causal to drive expression of immune-response like genes.

Our findings provide a missing link between metabolic rewiring, epigenetic modification, and aging in response to persistent DNA damage. Our results indicate that perturbation of any one of the three interconnected cellular processes-metabolism, epigenetic modification, and lipid oxidation is enough to rescue the deleterious effects of persistent DNA damage. Future studies could target any one of these pathways or collectively all these pathways to ameliorate chronic DNA damage associated health decline. Such strategies could include dietary supplementation of less oxidizable fatty acids such as MUFA, providing alternative acetyl-CoA substrates such as acetate or using pharmacological inhibitors of lipid oxidation.

## MATERIALS AND METHODS

### *C. elegans* maintenance and strains

C. elegans were grown at 20°C on nematode growth medium (NGM) agar plates seeded with *Escherichia coli* OP50 or HT115, unless specified. All strains were extensively backcrossed (∼5-10 times). Fresh stock was thawed every 3 months and PCR was used to confirm genotype. *E. coli* OP50 was cultured overnight in LB at 37°C. *E. coli* HT115 was used to perform RNAi and was cultured overnight at 37°C in LB containing ampicillin (100 μg/mL). Strains used in this study were listed in supplementary table 1.

### Oil Red O staining

Oil-Red-O staining was performed as described previously^30^. Briefly, ∼80-150 adult animals at indicated ages were washed three times with 1x PBS pH 7.4. To permeabilize the cuticle, worms were resuspended in 120 µl of PBS to which an equal volume of 2x MRWB buffer containing 2% paraformaldehyde (PFA) was added. 2x MRWB buffer: 160 mM KCl, 40 mM NaCl, 14 mM Na2EGTA, 1 mM spermidine-HCl, 0.4 mM spermine, 30 mM Na-PIPES pH 7.4, 0.2% ß-mercaptoethanol). Samples were gently rocked for 1h at room temperature. Animals were allowed to settle by gravity, buffer was aspirated, and worms were washed with 1x PBS to remove PFA. Worms were then resuspended in 60% isopropanol and incubated for 15 minutes at room temperature to dehydrate. Oil-Red-O is prepared as follows: a 0.5g/100mL isopropanol stock solution equilibrated for several days was freshly diluted to 60% with water and rocked for at least 1h, then filtered with 0.45 or 0.22µm-filter. After allowing worms to settle, isopropanol was removed, 120 µl of 60% Oil-Red-O stain was added, and animals were incubated overnight with rocking. Unbound dye was removed after allowing worms to settle, and adding 240 µL of 1x PBS 0.01% Triton X-100 was added. The worms were washed two times with 1x PBS pH 7.4. Animals were mounted on 2% agarose pads.

### Oil-Red-O quantification

Oil-Red-O worms were imaged at 10× magnification using an ECHO Revolve microscope and saved as TIF files. The same exposure settings were used across all conditions within each experiment. Using Cell Profiler processing software, raw images were inverted, separated in to RED, GREEN, and BLUE and thresholded to outline the worm body ^81^. The same threshold values were used across all conditions within each experiment. Oil red staining inside worms is then identified using a threshold value using inverted BLUE channel. The threshold values were uniform across all analysis. The area of the worm and the oil red o-stained area inside each worms were calculated. The data is represented as a fraction of oil red o-stained area to the total worm area. In order to normalize for experimental batch variations, each experiment carried control groups (L4440-N2) and all data are normalized to the average value of the control group. The data (a.u.) were graphed using Graphpad Prism, and statistical significance was determined two-way ANOVA with sidak multiple comparison test.

### Lipid analysis

Triacylglycerol levels were quantified using Biovision Kit for TAG assay Triacyl glyceride quantification kit as described with a few modifications^82^. Briefly, 2000 worms in the mentioned stage were collected washed with M9 buffer. The worms were then transferred to Precellys hard tissue homogenizing (2ml) tubes with 100 ul of water and lysed using 2 times for 20s lysis at 6800 rpm setting. After the lysis 100μL of 10% NP-40 alternative was added and the sample was heated to 100°C in water bath for 5 minutes and cooled down to room temperature. Boiling of the sample was repeated once again and the samples were centrifuged for 2 min at 1200 xg. The supernatants were collected and the TAG amounts were measured using the manufacture’s protocol.

Total lipids for PL analysis and Lipidomics were extracted using the folch lipid extraction protocol. Briefly, 2000 worms were lysed in 500 ul 0.75% KCL solution using the Precellys homogenizer. 3 ml of 2:1 chloroform methanol was added to the homogenized solution and incubated for 1hr with vigorous shaking at 4C. Samples were centrifuged at 3000 x g for 10 minutes at 4C for phase separation and organic layers were separated. To the aqueous and protein layers 2 ml of chloroform was added and the lipids were re-extracted. Both organic layers were pooled together dried under nitrogen and resuspended in 100 ul of 2:1 chloroform methanol containing 0.1% BHT. Total phospholipids were quantified using a micro method for phosphorus^83^.

### Quantitative RT–PCR

For experiments using whole worms, at least 500 age-synchronized worms in biological triplicate for each condition were harvested, washed three times with M9 and resuspended in 500□μl Trizol. To extract total RNA, worm or tissue pellets in Trizol underwent six freeze-thaw cycles in a dry ice-ethanol bath. RNA was extracted according to the Trizol procedure, and resuspended in RNase- and DNase-free water. RNA from whole worms was quantified using the Qubit 2.0 fluorometer (Invitrogen), and at least 500□ng total RNA per condition was used for cDNA synthesis^37^. Total RNA was then reverse-transcribed using Transcriptor First Strand cDNA Synthesis Kit (Roche), based on the manufacturer’s instructions. RT–qPCR was performed using diluted cDNA on a Quantsudio3 (applied biosystems) with PowerUp SYBR green (Invitrogen). Primers used in RT–qPCR are listed in supplementary table 2. Melt curves were examined to ensure primer specificity. Results were analyzed using the standard ΔΔ*C*T method and values were normalized to *rpl-32* or *act-1* as an internal control. All data shown represent at least three biologically independent samples.

### Lifespan analysis

Worms were age-synchronized by egg-lay. For all experiments, every genotype and condition was performed in parallel at 20°C. Worms were transferred to RNAi plates as D1 (day 1) adults. To prevent progeny production, during adulthood, worms were transferred to 2.5 μgml-1 FUDR-containing NGM seeded plates every two days. 60–100 animals were assayed for each condition/genotype. Worms were scored every second day. Death was indicated by total cessation of movement in response to gentle mechanical stimulation^24^. Prism6 software was used for statistical analysis. Log-rank (Mantel-Cox) method was used to determine the significance difference.

### Intestinal nuclei analysis

Dissection and fixation were carried out by freeze-crack method^84^. Worms were dissected in M9 buffer on a polylysine-coated slide. The slides were fixed in ice cold ethanol for 2 min. Next, the slides were placed in phosphate-buffered saline (PBS) and washed with fresh PBS for 5 min. Slides were then mounted with 4′,6-diamidino-2-phenylindol (DAPI) solution (VECTASHIELD). We used at least 30 worms in each sample for DAPI staining and intestinal nuclei from more than 10 worms were selected and quantified for nuclear size ^84^.

### Image analysis of intestinal nuclei

Intestinal cells were imaged at ×63 magnification using Leica-TCS SP8, laser scanning confocal microscope. Z-stacks of the intestine were analyzed using LAS X. Contours of individual nuclei, according to DAPI fluorescence, were manually defined and used to render nuclei area. Maximum cross-section area of each nucleus was acquired from the surface tool statistics by Image J.

### Western blot analysis

Age-synchronized worms ∼1500 were washed off from the plate with M9 buffer and frozen in liquid nitrogen. Equal volume of sample buffer (0.1 M Tris pH 6.8, 7.5 M urea, 2 % SDS, 100 mM β-ME, 0.05 % bromophenol blue) was added to worms Worm extracts were generated by sonication by heating worms to 65 °C for 10 min, sonicating for two 30-s bursts, heating to 95 °C for 5 min and then kept at 37 °C until loading onto SDS-PAGE gel^85^. Proteins were transferred to nitrocellulose and blotted with the following antibodies: H4K8/16ac (active motif 39930) at 1:1000, H3K9ac (CST, C5B11) at 1:1000, and β-actin (Sigma, A3854) at 1:1000.

### Compound treatment assay

*Pyruvate, Acetate, Citrate treatment*. We performed metabolites treatment assays by adding pyruvate, acetate, citrate to a final concentration of 50mM on the NGM plates^84^. Worms were transferred on compound plates with FUDR as day 1 adults and transferred every day until D7.

### Transcriptomics

*Sample Preparation*: At least 500 age-synchronized worms in biological triplicate for each condition were harvested, washed five times with M9 and resuspended in 500□μl Trizol. RNA extraction was carried out as mentioned above. *RNA quantification and qualification: RNA* degradation and contamination was monitored on 1% agarose gels. RNA purity was checked using the NanoPhotometer®spectrophotometer (IMPLEN, CA, USA). RNA integrity and quantitation were assessed using the RNA Nano 6000 Assay Kit of the Bioanalyzer 2100 system (Agilent Technologies, CA, USA).

*Library preparation for Transcriptome sequencing:* A total amount of 1 μg RNA per sample was used as input material for the RNA sample preparations. Sequencing libraries were generated using NEBNext®Ultra™RNA Library Prep Kit for Illumina® (NEB, USA) following manufacturer’s recommendations and index codes were added to attribute sequences to each sample. Briefly, mRNA was purified from total RNA using poly-T oligo-attached magnetic beads. Fragmentation was carried out using divalent cations under elevated temperature in NEB Next First Strand Synthesis Reaction Buffer (5X). First strand cDNA was synthesized using random hexamer primer and M-MuLV Reverse Transcriptase (RNase H-). Second strand cDNA synthesis was subsequently performed using DNA Polymerase I and RNase H. Remaining overhangs were converted into blunt ends via exonuclease/polymerase activities. After adenylation of 3’ ends of DNA fragments, NEB Next Adaptor with hairpin loop structure were ligated to prepare for hybridization. In order to select cDNA fragments of preferentially 150∼200 bp in length, the library fragments were purified with AMPure XP system (Beckman Coulter, Beverly, USA). Then 3 μL USER Enzyme (NEB, USA) was used with size-selected, adaptor-ligated cDNA at 37° C for 15 min followed by 5 min at 95 °C before PCR. Then PCR was performed with Phusion High-Fidelity DNA polymerase, Universal PCR primers and Index (X) Primer. At last, PCR products were purified (AMPure XP system) and library quality was assessed on the Agilent Bioanalyzer 2100 system.

#### Clustering and sequencing

The clustering of the index-coded samples was performed on a cBot Cluster Generation System using PE Cluster Kit cBot-HS (Illumina) according to the manufacturer’s instructions. After cluster generation, the library preparations were sequenced on an Illumina platform and 125 bp/150 bp paired-end reads were generated.

#### Quality control

Raw data (raw reads) of fastq format were firstly processed. In this step, clean data (clean reads) were obtained by removing reads containing adapter, reads containing ploy-N and low-quality reads from raw data. At the same time, Q20, Q30 and GC content the clean data were calculated. All the downstream analyses were based on the clean data with high quality.

Reads mapping to the reference genome: Reference genome and gene model annotation files were downloaded from genome website directly. Index of the reference genome was built using hisat2 2.1.0 and paired-end clean reads were aligned to the reference genome using HISAT2. We selected HISAT2 as the mapping tool for that Hisat2 can generate a database of splice junctions based on the gene model annotation file and thus a better mapping result than other non-splice mapping tools, which is a fast and sensitive alignment program for mapping next-generation sequencing reads against the general human population.

Quantification of gene expression level: Feature Counts v1.5.0-p3 was used to count the reads numbers mapped to each gene^86^. Then the FPKM of each gene was calculated based on the length of the gene and reads count mapped to this gene. FPKM, expected number of Fragments Per Kilobase of transcript sequence per Millions base pairs sequenced, considers the effect of sequencing depth and gene length for the reads count at the same time, and is currently the most commonly used method for estimating gene expression levels^87^.

#### Differential expression analysis

(For DESeq2 with biological replicates) Differential expression analysis between two conditions/groups (two biological replicates per condition) was performed using the DESeq2 R package (1.14.1). DESeq2 provide statistical routines for determining differential expression in digital gene expression data using a model based on the negative binomial distribution. The resulting P-values were adjusted using the Benjamini and Hochberg’s approach for controlling the False Discovery Rate (FDR). Genes with an adjusted P-value <0.05 found by DESeq2 were assigned as differentially expressed. (For edgeR without biological replicates) Prior to differential gene expression analysis, for each sequenced library, the read counts were adjusted by edgeR program package through one scaling normalized factor. Differential expression analysis of two conditions was performed using the edgeR R package (3.16.5). The P values were adjusted using the Benjamini & Hochberg method. Corrected P-value of 0.05 and absolute foldchange of 1 were set as the threshold for significantly differential expression. Pathway enrichment analysis was performed using Wormcat online tool as described in Holdorf el al ^27^. Figures from the data was made using ggplot2 or R language.

### Metabolomics

#### Sample Preparation

A Synchronous population of 500 young adult worms were washed off the plates in M9 buffer and collected in 1.5mL Eppendorf tubes. The worms were further washed, three times with 0.5mL of mass-spec grade water. After the final wash, the worms were collected in 0.5mL ice-cold 80% methanol into 2mL-homogenizer tubes. The worms were homogenized with a 5□mm steel bead using a TissueLyser II (Qiagen) for 2□×□2.5□min at a frequency of 30 times/sec, followed by a tip sonication (energy level: 40 joules; output: 8 watts) for two times on ice water. Samples were centrifuged at 14,000 rpm and 4 °C for 10 min, the supernatant was transferred to a fresh Eppendorf tube and the solvent was removed. The residue was dissolved in 100 μl MeOH and transferred to an autosampler vial following centrifugation to remove any particulate matter.

### Untargeted high-resolution LC-HRMS

#### Sample preparation

Metabolic quenching and polar metabolite pool extraction was performed by adding ice cold 80% methanol at a ratio of 1:15 weight:volume. Internal standards; deuterated (D_3_)-creatinine and (D_3_)-alanine, (D_4_)-taurine, (D_3_)-lactate (Sigma-Aldrich) at a final concentration of 10µM. Samples were homogenized for 40 seconds at 60hz using an MP Bio FastPrep with Matrix A beads. The supernatant was cleared of protein by centrifugation at 16,000xg. 2µL of cleared supernatant was subjected to online LC-MS analysis.

### LC-HRMS Method (Polar)

Analyses were performed by untargeted LC-HRMS. Briefly, Samples were injected via a Thermo Vanquish UHPLC and separated over a reversed phase Thermo HyperCarb porous graphite column (2.1×100mm, 3μm particle size) maintained at 55°C. For the 20 minute LC gradient, the mobile phase consisted of the following: solvent A (water / 0.1% FA) and solvent B (ACN / 0.1% FA). The gradient was the following: 0-1min 1% B, increase to 15%B over 5 minutes, continue increasing to 98%B over 5 minutes, hold at 98%B for five minutes, reequillibrate at 1%B for five minutes. The Thermo IDX tribrid mass spectrometer was operated in both positive and ion mode, scanning in ddMS^2^ mode (2 μscans) from 100 to 800 m/z at 70,000 resolution with an AGC target of 2e5 for full scan, 2e4 for ms^2^ scans. Source ionization setting was 3.0 and 2.4 kV spray voltage, respectively, for positive and negative mode. Source gas parameters were 35 sheath gas, 12 auxiliary gas at 320°C, and 8 sweep gas. Calibration was performed prior to analysis using the Pierce™ FlexMix Ion Calibration Solutions (Thermo Fisher Scientific). Integrated peak areas of known identity from in-house libraires were then extracted manually using Quan Browser (Thermo Fisher Xcalibur ver. 2.7). Untargeted differential comparisons were performed using Compound Discoverer 3.0 (Thermo Fisher) to generate a ranked list of significant compounds with tentative identifications from BioCyc, KEGG, and internal databases.

#### Data Analysis

Levels of TCA metabolites, carnitine and acetyl-carnitine, and AA were quantifies using Quan browser software. Un targeted analysis of metabolomics data was performed using compound discoverer software. Peaks with 10000 ion intensity and 10 S/N was identified and various adducts are consolidated using m/z and the retention time. compounds are then queried against Pubchem and Human metabolomic database and MZ cloud fragmentation database to obtain putative identifications. Peaks with more than one identity is confirmed based on manual fragmentation analysis. Data was then normalized based on the internal standard analyzed using Graph pad prism software. PCA analysis was performed using SIMCA software

### Transcriptomic and Metabolomics data integration

Enriched transcriptomic and metabolomics data was used for transcriptomic data integration. List of enriched transcripts and metabolites were submitted to the MOFA2 software in R language^29^. A Model was developed with 2 factors. Both the metabolites and the transcript with the normalized future weight >1 was selected as the transcripts/metabolites with highly coordinated levels. These transcripts were then subjected to pathway enrichment analysis using wormcat as described previously ^27^.

### Acetyl-CoA measurements

2000 worms were washed in 1X Tris buffered saline at least 3 times to get rid of bacteria. Wash the worms with sterile water 2 times and then add 1mL of 10% ice-cold TCA (TCA solution in - 20C). Collect in percelly’s tubes. Homogenize “Hard” 2 times and collect the lysates in pre-chilled 1.5mL eppendorf tubes and use for acetyl-coA quantitation.

Acetyl-CoA was measured by liquid chromatography-high resolution mass spectrometry as previously reported^88^. Briefly, samples and calibrators generated from commercially available standards (from Sigma-Aldrich) were spiked with a constant amount of ^13^C_3_^15^N_1_-acyl-CoA extract as an internal standard generated biosynthetically as previously described from ^13^C_3_^15^N_1_-pantothenate^89^. Samples were sonicated, then extracted by Oasis HLB solid phase extraction, evaporated to dryness, resuspended in 50 µL 5% 5-sulfosalicylic acid in water and 10 µL was injected for analysis on a Ultimate 3000 UHPLC (Thermo) with a Waters HSS T3 column (2.1 × 100 mm) coupled to a Q Exactive Plus (Thermo) operating in positive ion mode alternating full scan from 760–1800 m/z at 140,000 resolution and data independent acquisition (DIA) looped 3 times with all fragment ions multiplexed at a normalized collision energy (NCE) of 20 at a resolution of 280,000. Operating conditions on the mass spectrometer were as follows; auxiliary gas 10 arbitrary units (arb), sheath gas 35 arb, sweep gas 2 arb, spray voltage 4.5 kV, capillary temperature 425 °C, S-lens RF-level 50, aux gas heater temperature 400°C, in-source CID 5 eV. LC conditions were column oven temperature 30°C, solvent A water with 5 mM ammonium acetate, solvent B 95:5 acetonitrile: water with 5 mM ammonium acetate, solvent C (wash solvent) 80:20 acetonitrile: water with 0.1% formic acid with the following gradient of 0.2 mL/min flow at 98% A and 2% B for 1.5 min, 80% A 20% B at 5 min, 100% B at 12 min, 0.3 mL/min 100% B at 16 min, 0.2 mL/min 100% C at 17 min, held to 21 min, then re-equilibrated at 0.2 mL/min flow at 98% A and 2% B from 22 to 28 min. Flow from 4–18 minutes was diverted to the instrument. Data was processed in Xcalibur and TraceFinder 4.1 (Thermo). Protein normalization was by a Pierce BCA kit according to manufactures instructions.

### Lipidomics

Lipidomics analysis was performed using liquid chromatography high resolution tandem mass spectrometry. Samples were analyzed using Dionex Ultimate 3000 HPLC system coupled on-line to a Q-Exactive Hybrid Quadrupole-Orbitrap Mass Spectrometer (Thermo Fisher) A reverse phase, C-30 column phase column (Accucore™ C30, 2.6 µm, 150 Å, 250 × 2.1 mm, thermo Scientific™) and a binary gradient consisting of Solvent A (1:1 acetonitrile:water with 0.1% formic acid and 5 mM ammonium formate) and Solvent B(8.5:1:0.5 2-Propanol:Acetonitrile:water with 0.1% formic acid and 5 mM ammonium formate) was used for the chromatographic separation. Five nano moles of phospholipid samples were dried and dissolved in solvent B and injected in the LC using the autosampler. The following elution gradient was used: 0-20 minutes: an isocratic flow of 30% solvent B at 100 µL per minute, 20-50 minutes: a liner gradient of 30% to 100 % solvent B at a flow rate of 100 µL per minute, 50-70 minutes: an isocratic flow of 100% Solvent B at 100 µL per minute, 70-85 minutes: linear gradient of 100 % to 30% solvent B at 100 µL per minute followed by 10 minutes re-equilibration of column at 30% solvent B at 100 µL per minute. Sample from LC was injected into the mass spectrometer using a Heated electron spray ionization (HESI) source. For phospholipid analysis mass spectrometer was operated at negative ion mode with spray voltage of 2.5 KV, capillary temperature of 320°C, sheath gas 6 AU, S-lens RF 60 mass range 150-1800 m/z with a resolution of 70, 000 FWHM at 200 m/z. Tandem mass spectrometry was performed on Top 10 high ions using data dependent mode. Higher energy collisional dissociation (HCD) method with 24% energy was used. For Triacyl glycerol analysis, the mass spectrometer was operated in a positive ion mode with the spray voltage of 3.5 KV. Data was analyzed using Compound discoverer software as described previously^90^. Identity of the lipids were confirmed by the representative fragmentation analysis. Statistical analysis was performed using graph pad prism software.

## Acknowledgements

We thank the *C. elegans* Genetics Center for strains. We thank J. Yanowitz, A. Ghazi, T. Lamintina and M. Gill for RNAi and for insightful scientific discussions; T. Finkel for edits and comments to the manuscript. This work was funded by NIH grants R00AG049126 (AUG), S10OD023402 (SGW), R01GM132261 (NWS), AI145406, CA165065,CA243142,AI068021, GM113908, HL114453, AI156924, NS076511, AI15 6923, NS061817 (H.B./ V.K.)

## Author contributions

AUG and SH provided study design. SH and AUG performed most of the experiments. S. Han, HS, ES and AS provided technical assistance and blinded ORO imaging. SH, TA, SJM and SGW performed metabolomics and analysis. SH, TA, HB and VK performed lipidomics and analysis. HLP and NWS performed acetyl-CoA measurements. PS helped with intellectual and technical expertise on epigenetic analysis. SH and AUG drafted the manuscript with the help of all co-authors. AUG supervised the study.

## Notes

### Competing Interest Statement

The authors have declared no competing interest.

