## Supplementary material for "Persistent DNA damage rewires lipid metabolism and promotes histone hyperacetylation via MYS-1/Tip60": Supp Figures

### Slide 1
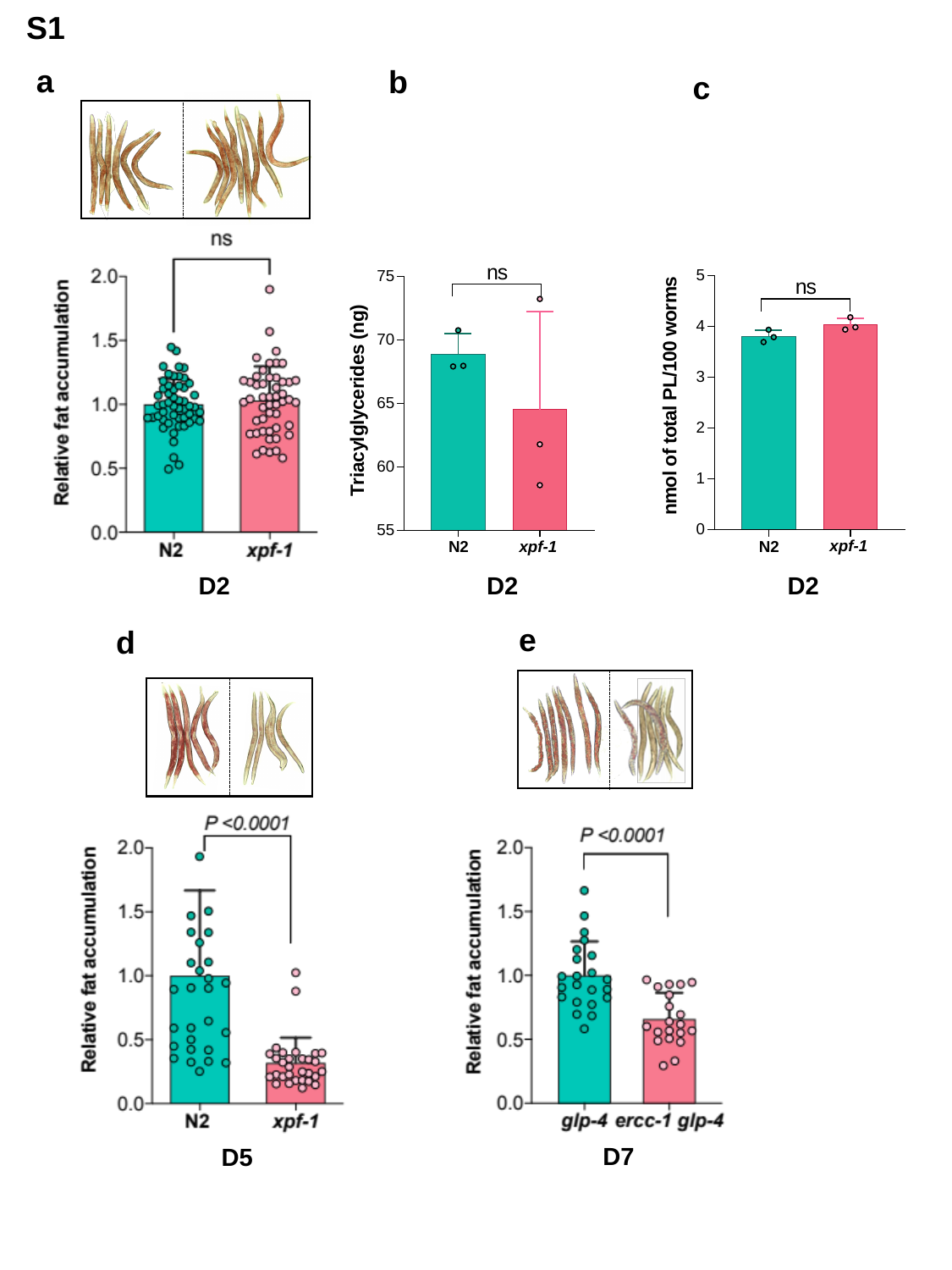

S1
a
b
c
D2
D2
D2
e
d
D7
D5

### Slide 2
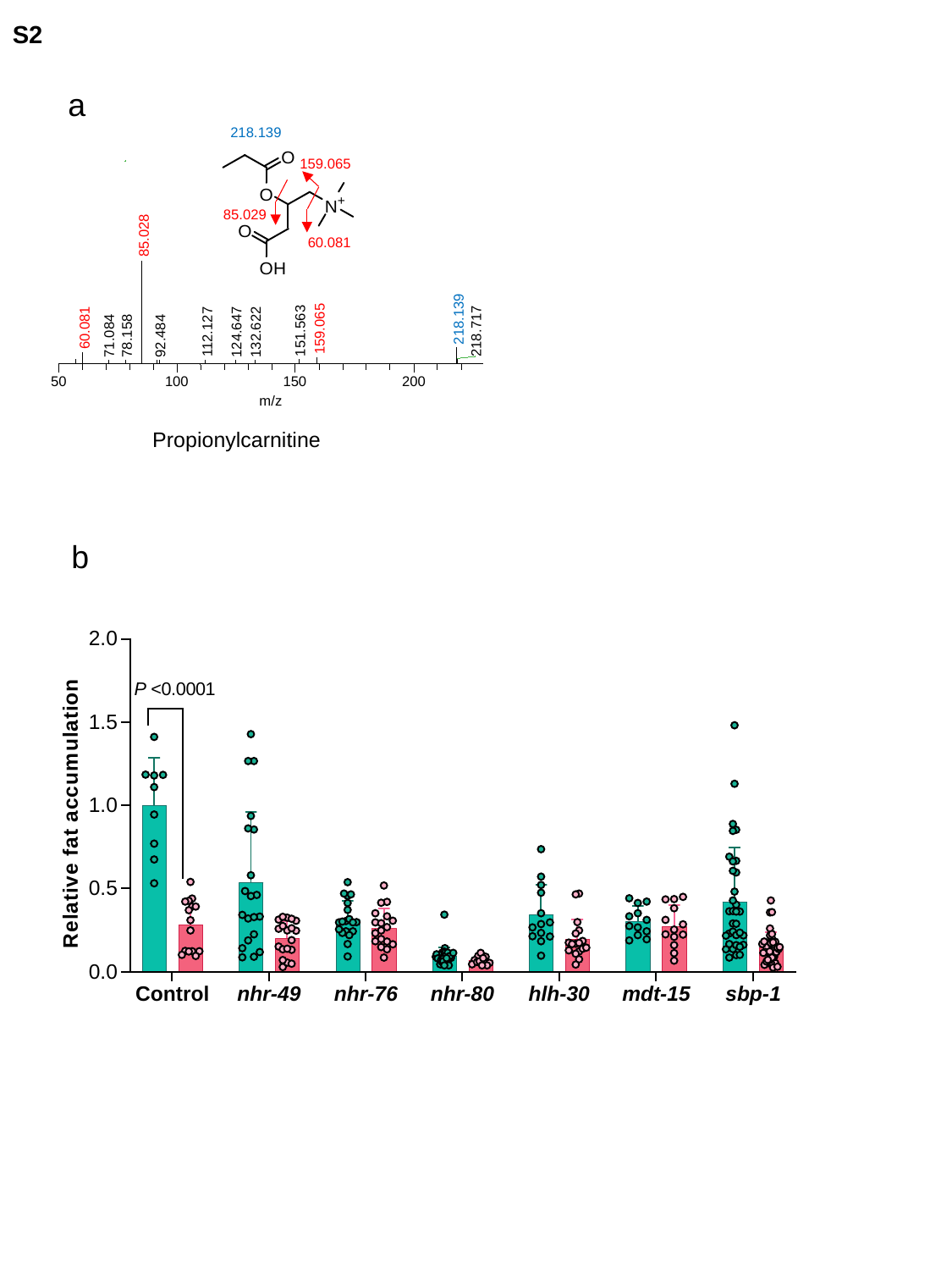

S2
a
218.139
159.065
85.029
85.028
218.139
60.081
159.065
151.563
218.717
112.127
124.647
132.622
71.084
78.158
92.484
50
100
150
200
m/z
60.081
Propionylcarnitine
b

### Slide 3
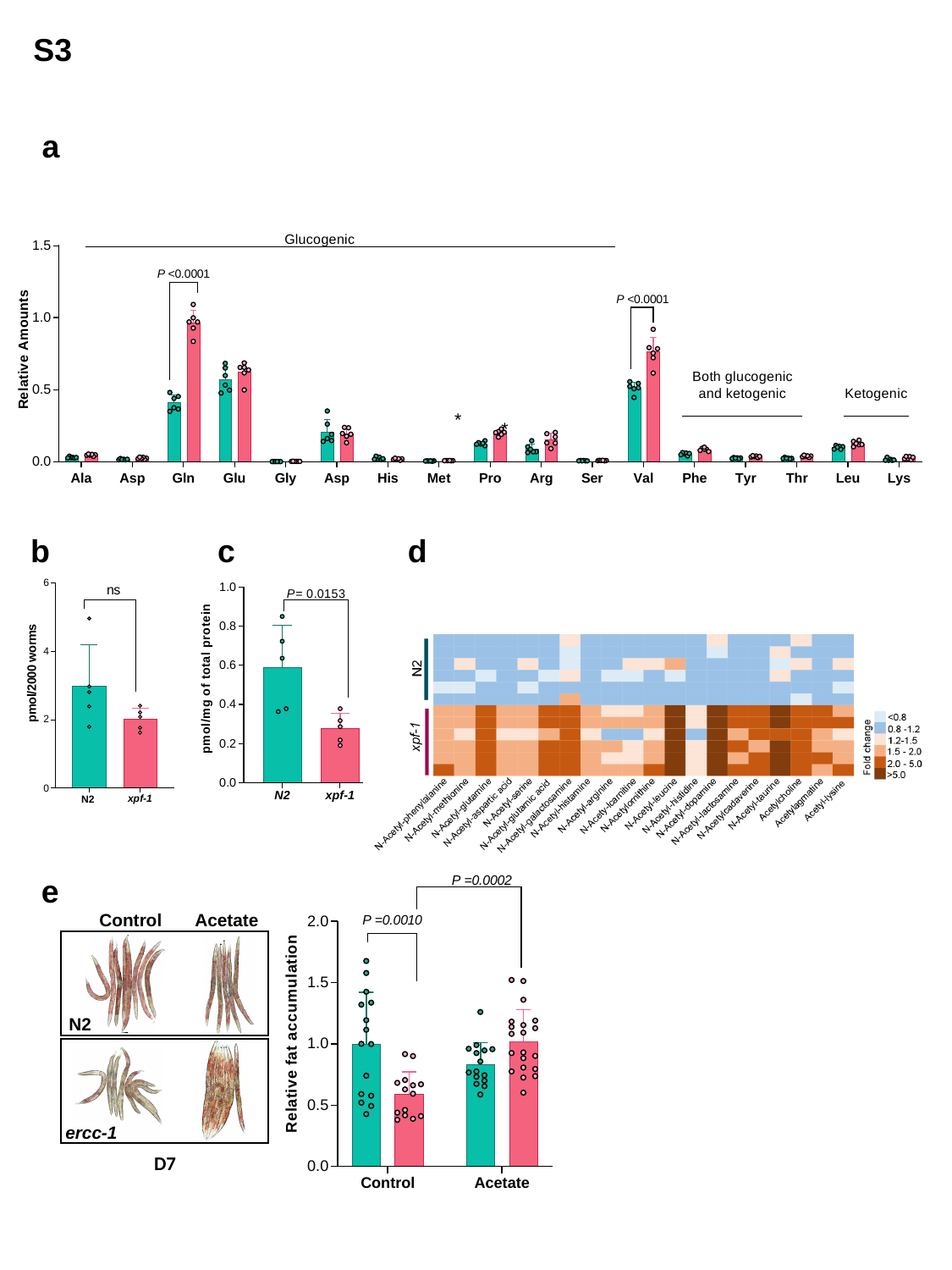

S3
a
b
c
d
e
Control
Acetate
N2
ercc-1
D7

### Slide 4
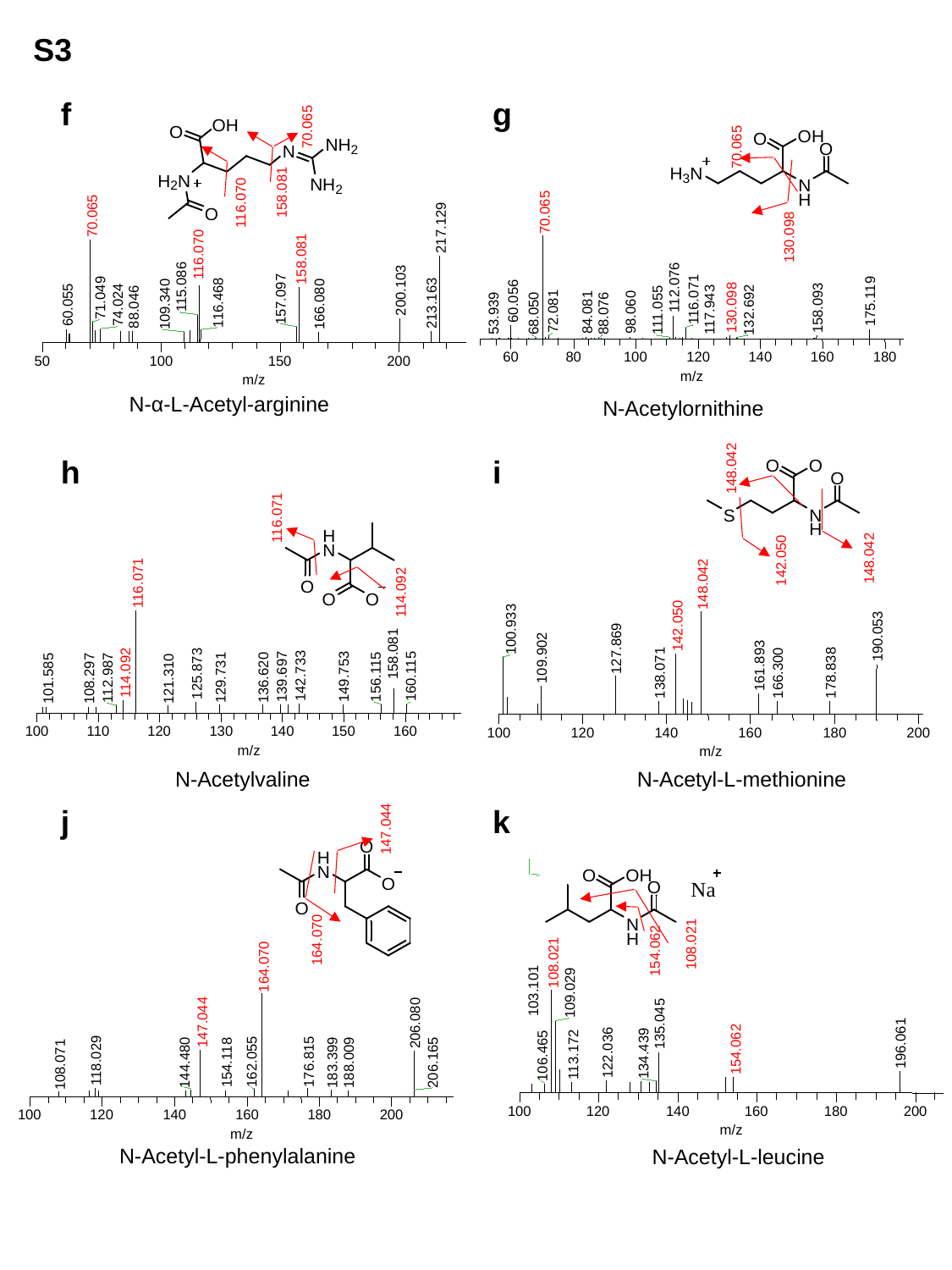

S3
f
g
70.065
70.065
70.065
217.129
116.070
158.081
115.086
200.103
71.049
157.097
116.468
213.163
109.340
166.080
60.055
74.024
88.046
50
100
150
200
m/z
158.081
70.065
112.076
116.071
60.056
175.119
158.093
130.098
111.055
117.943
132.692
72.081
84.081
98.060
53.939
68.050
88.076
60
80
100
120
140
160
180
m/z
116.070
130.098
N-α-L-Acetyl-arginine
N-Acetylornithine
h
i
148.042
116.071
114.092
148.042
142.050
116.071
158.081
125.873
114.092
142.733
160.115
139.697
149.753
156.115
112.987
129.731
136.620
101.585
108.297
121.310
100
110
120
130
140
150
160
m/z
148.042
142.050
100.933
190.053
127.869
109.902
161.893
138.071
166.300
178.838
100
120
140
160
180
200
m/z
N-Acetyl-L-methionine
N-Acetylvaline
j
k
147.044
108.021
103.101
109.029
135.045
196.061
154.062
122.036
134.439
113.172
106.465
100
120
140
160
180
200
m/z
164.070
108.021
164.070
206.080
147.044
118.029
162.055
176.815
144.480
154.118
183.399
188.009
206.165
108.071
100
120
140
160
180
200
m/z
154.062
N-Acetyl-L-phenylalanine
N-Acetyl-L-leucine

### Slide 5
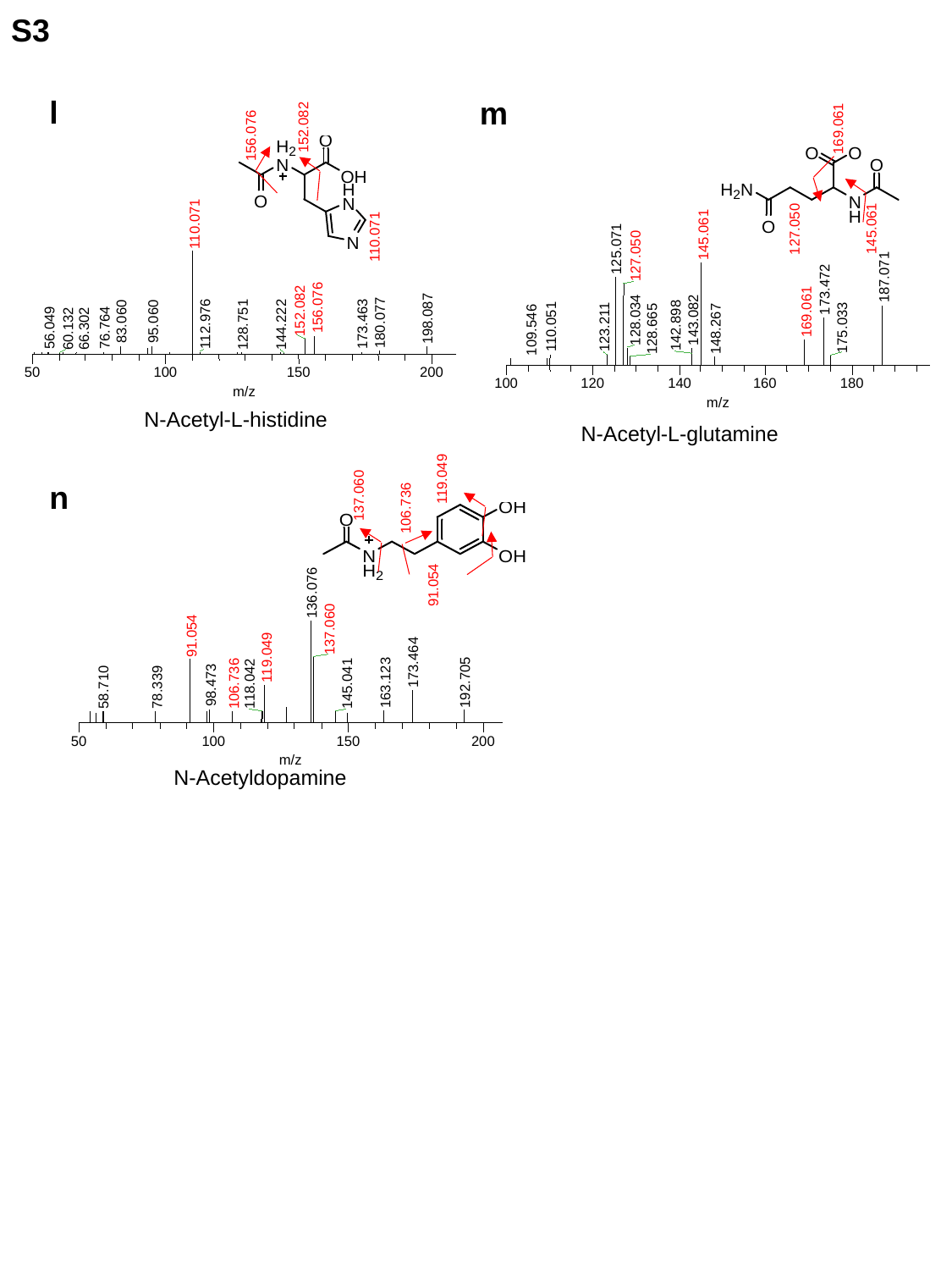

S3
l
m
152.082
156.076
110.071
156.076
152.082
198.087
83.060
95.060
180.077
112.976
173.463
128.751
144.222
56.049
76.764
60.132
66.302
50
100
150
200
m/z
110.071
169.061
145.061
125.071
127.050
187.071
173.472
169.061
128.034
143.082
142.898
110.051
123.211
175.033
128.665
148.267
109.546
100
120
140
160
180
m/z
145.061
127.050
N-Acetyl-L-histidine
N-Acetyl-L-glutamine
119.049
137.060
106.736
91.054
n
136.076
137.060
91.054
119.049
173.464
192.705
163.123
118.042
145.041
106.736
98.473
58.710
78.339
50
100
150
200
m/z
N-Acetyldopamine

### Slide 6
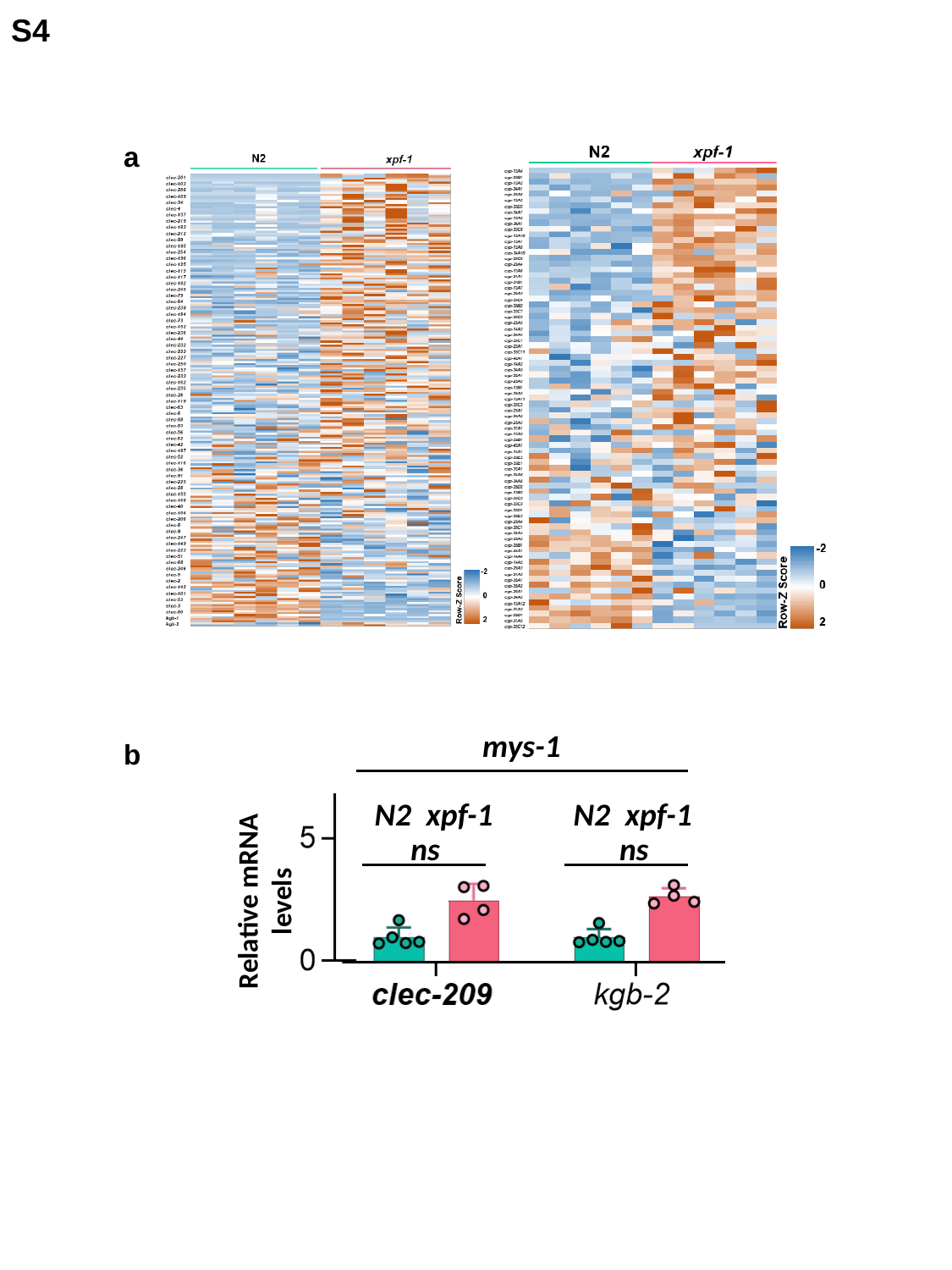

S4
a
mys-1
b
N2
xpf-1
N2
xpf-1
ns
ns
Relative mRNA levels

### Slide 7
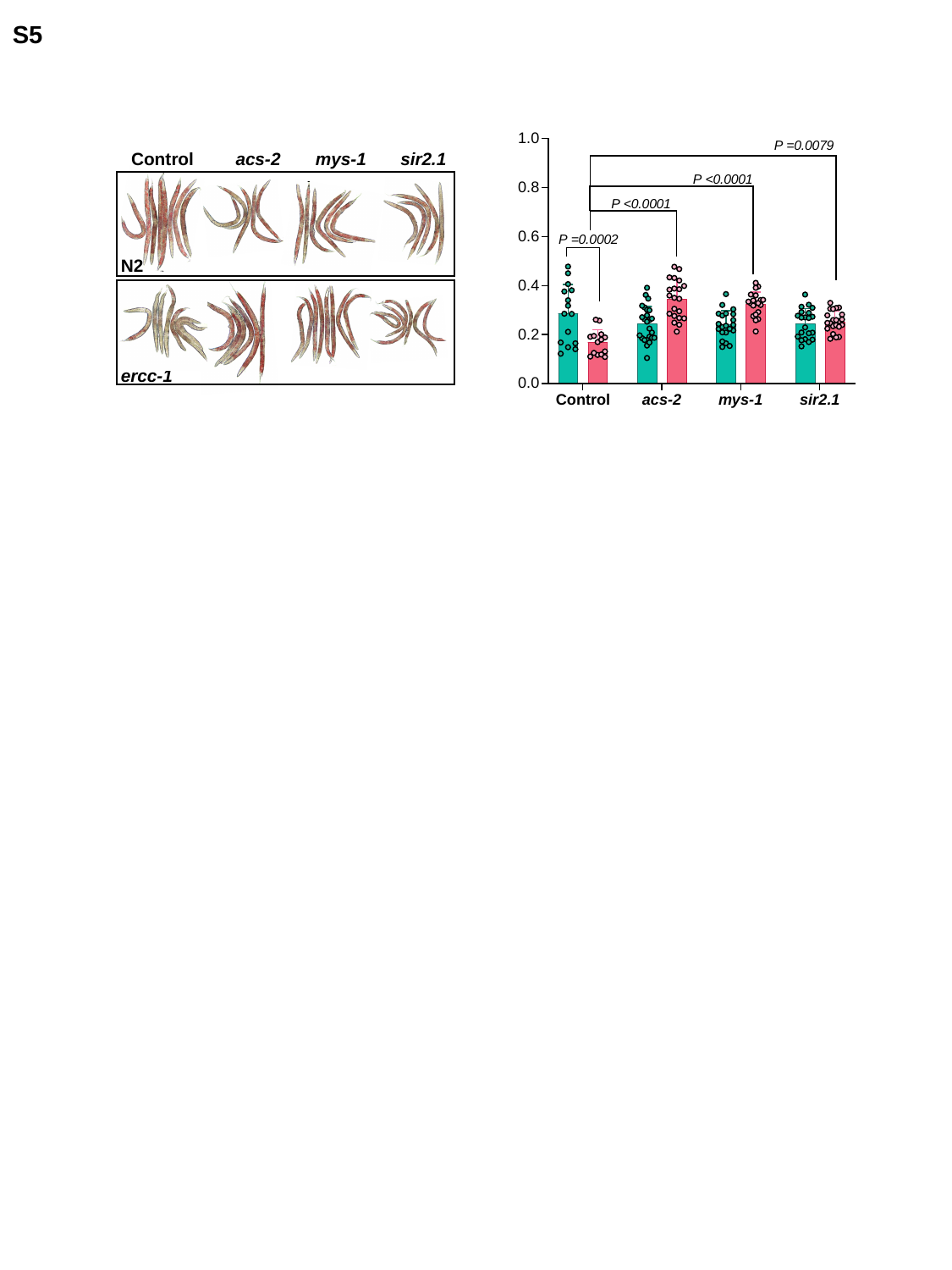

S5
Control
acs-2
mys-1
sir2.1
N2
ercc-1

### Slide 8
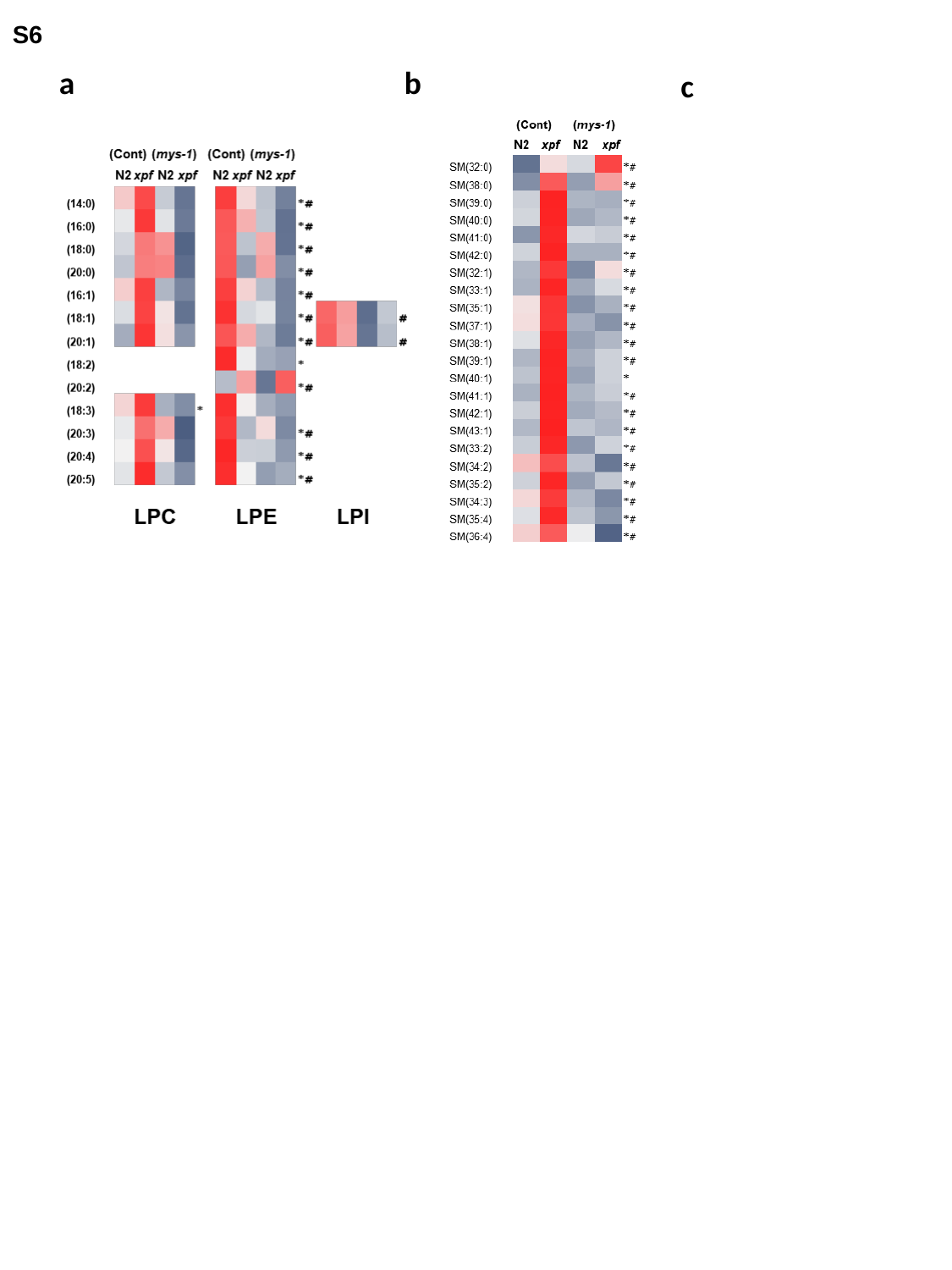

S6
a
b
c
